# Click-PD: A Quantitative Method for Base-Modified Aptamer Discovery

**DOI:** 10.1101/626572

**Authors:** Chelsea K. L. Gordon, Diana Wu, Trevor A. Feagin, Anusha Pusuluri, Andrew T. Csordas, Michael Eisenstein, Craig J. Hawker, Jia Niu, H. Tom Soh

## Abstract

Base-modified aptamers that incorporate non-natural chemical moieties can achieve greatly improved affinity and specificity relative to natural DNA or RNA aptamers. However, conventional methods for generating base-modified aptamers require considerable expertise and resources. In this work, we have accelerated and generalized the process of generating base-modified aptamers by combining a click-chemistry strategy with a fluorescence-activated cell sorting (FACS)-based screening methodology that measures the affinity and specificity of individual aptamers at a throughput of ∼10^7 per hour. Our “click-PD” strategy offers many advantages. First, almost any chemical modification can be introduced with a commercially-available polymerase. Second, click-PD can screen vast numbers of individual aptamers based on quantitative on- and off-target binding measurements to simultaneously achieve high affinity and specificity. Finally, it requires minimal specialized equipment or reagents besides a FACS instrument which is now widely-available. Using click-PD, we generated a boronic acid-modified aptamer with ∼1 μM affinity for epinephrine, a target for which no aptamer has been reported to date. We subsequently generated a mannose-modified aptamer with nanomolar affinity for the lectin concanavalin A (ConA). The strong affinity of both aptamers is fundamentally dependent upon the presence of chemical modifications, and we show that their removal essentially eliminates aptamer binding. Importantly, our ConA aptamer exhibited exceptional specificity, with minimal binding to other structurally-similar lectins. Finally, we show that our aptamer remarkable biological activity. Indeed, to the best of our knowledge, this aptamer is the most potent inhibitor of Con A-mediated hemagglutination reported to date.

## INTRODUCTION

Base-modified aptamers that incorporate non-natural chemical functional groups provide many advantages as affinity reagents because, like conventional aptamers, they are chemically synthesized and sequence-defined, while also offering a much broader chemical repertoire than their natural DNA and RNA counterparts^1^. This can result in a commensurate expansion in the spectrum of targets that can be recognized with high affinity and specificity, and a number of groups have developed effective strategies for introducing such modifications into nucleic acids over the past two decades.^2–5^ For example, Gold and others developed a pioneering strategy for incorporating numerous modifications into nucleotide side-chains to produce base-modified aptamers that tightly bind to a wide range of biomolecules that otherwise do not interact with natural DNA or RNA.^3,6^ More recently, other groups have modified the backbones of nucleic acids to generate ‘xenobiotic’ nucleic acid (XNA) aptamers with novel chemical functionalities^7^, including exceptional stability in complex biological milieus.^8,9^

The process of generating base-modified aptamers typically entails the use of engineered polymerases that can faithfully incorporate and amplify both natural and chemically-modified nucleotides while maintaining minimal error rates.^10^ Unfortunately, in addition to the considerable time and resources associated with engineering such polymerases, the resulting enzymes may not exhibit sufficient processivity or fidelity for efficient aptamer selection,^11,12^ and each new chemical modification may require a new campaign of polymerase engineering. Furthermore, the structural and biochemical requirements of a functional polymerase impose severe constraints on the extent of modification that is feasible,^11^ and nucleotides bearing especially large chemical modifications simply cannot be tolerated by any polymerase.

We have developed a high-throughput screening strategy that offers a general framework for the rapid and efficient selection of base-modified aptamers bearing virtually any chemical modification without the need for specially engineered polymerases. Our method builds on the click-SELEX technique described by the Krauss^13,14^ and Mayer groups,^15^ in which natural DNA nucleotides are substituted with an alkyne-modified alternative that can then be readily coupled to an azide-modified functional group. Whereas these previous methods relied on conventional SELEX, we have combined the click chemistry approach with the particle display (PD) platform previously described by our laboratory.^16,17^

‘Click-PD’ offers the critical capability to measure the affinity and specificity of every base-modified aptamer in a large library. Based on this measurement, each aptamer is then sorted individually using a fluorescence-activated cell sorter (FACS). To achieve this, click-PD uses a commercially-available polymerase to introduce nucleobases with chemical modifications into a natural DNA library, which has been converted into a pool of monoclonal ‘aptamer particles’ via an emulsion PCR process. These modified bases are then subjected to a click-chemistry reaction to enable covalent linkage of virtually any chemical group of interest. We demonstrate here that this approach is compatible with the copper(I)-catalyzed alkyne-azide cycloaddition (CuAAC) reaction as well as a copper-free click reaction^18^ based on strain-promoted alkyne-azide cycloaddition (SPAAC)^19,20,21^ The resulting libraries of base-modified aptamer particles are then incubated with target and non-target molecules that have been differentially labeled with distinct fluorophores. Finally, FACS is employed to isolate individual base-modified aptamer sequences that exhibit both high affinity and specificity for their intended target.

To demonstrate the generality of the method, we used click-PD to generate base-modified aptamers that incorporate distinct modifications that were specifically chosen to improve their binding performance for two very different target molecules. First, we used click-PD to efficiently screen a library of boronic acid-modified aptamers against the small-molecule target epinephrine, identifying a novel aptamer with an equilibrium dissociation constant (*K*_*d*_) of ∼1.1 μM. Next, we performed a multicolor click-PD screen to isolate mannose-modified aptamers that exhibit low nanomolar affinity and remarkable specificity for the lectin concanavalin A (Con A), with minimal binding to other lectins. This is especially notable as past efforts to select affinity reagents for lectins have tended to result in molecules with poor specificity. Importantly, one of our Con A aptamers also showed exceptional biological activity; to the best of our knowledge, it is the most potent inhibitor of Con A-mediated hemagglutination reported to date.^22^ Click-PD requires no specialized instrumentation other than a simple emulsifier and a FACS instrument, which is now available in many research settings. We therefore believe that click-PD will enable researchers to efficiently generate custom affinity reagents that have been modified with a wide range of chemical functionalities for a diverse variety of biological and biomedical applications.

## RESULTS

### Overview of the click-PD process

Our click-PD screening platform (**Fig. 1**) employs a click chemistry reaction to generate large libraries of monoclonal aptamer particles that each display many copies of a single base-modified aptamer sequence on their surface. These particles are then incubated with fluorescently-labeled target molecules, after which the affinity and specificity of the base-modified aptamer particles are individually measured via FACS. Those exceeding user-defined thresholds are collected for further selection or analysis.^16,17^ For this study, we employed nucleic acid libraries comprising a 40-nt random region flanked by primer-binding sites at both ends (see **Table S1** for all sequences). This library is first subjected to ‘pre-enrichment’ with bead-immobilized target molecules in order to reduce the size of the library from ∼10^14^ sequences to a scale that can be readily interrogated using FACS without reducing overall diversity (see Methods). We then convert the pre-enriched aptamers into a library of particles displaying aptamers containing DBCO-dUTP (for SPAAC) or C8-alkyne-dUTP (for CuAAC) via emulsion PCR (**Fig. 1**, step 1). We specifically selected DBCO-dUTP and C8-alkyne-dUTP because these nucleotide analogues can be incorporated by commercial DNA polymerases during the PCR procedure.^23–27^ We prepared water-in-oil emulsions with forward primer (FP)-coated magnetic beads, aptamer templates, and PCR reagents including both natural and modified deoxynucleotide triphosphates (dNTPs) under conditions such that each droplet contains (in most cases) one DNA template and one bead. Emulsion PCR amplification yields a library of monoclonal particles displaying sequences that bear alkyne groups (step 2). Details of this process are provided in the Supplementary Methods section.

**Figure 1.**
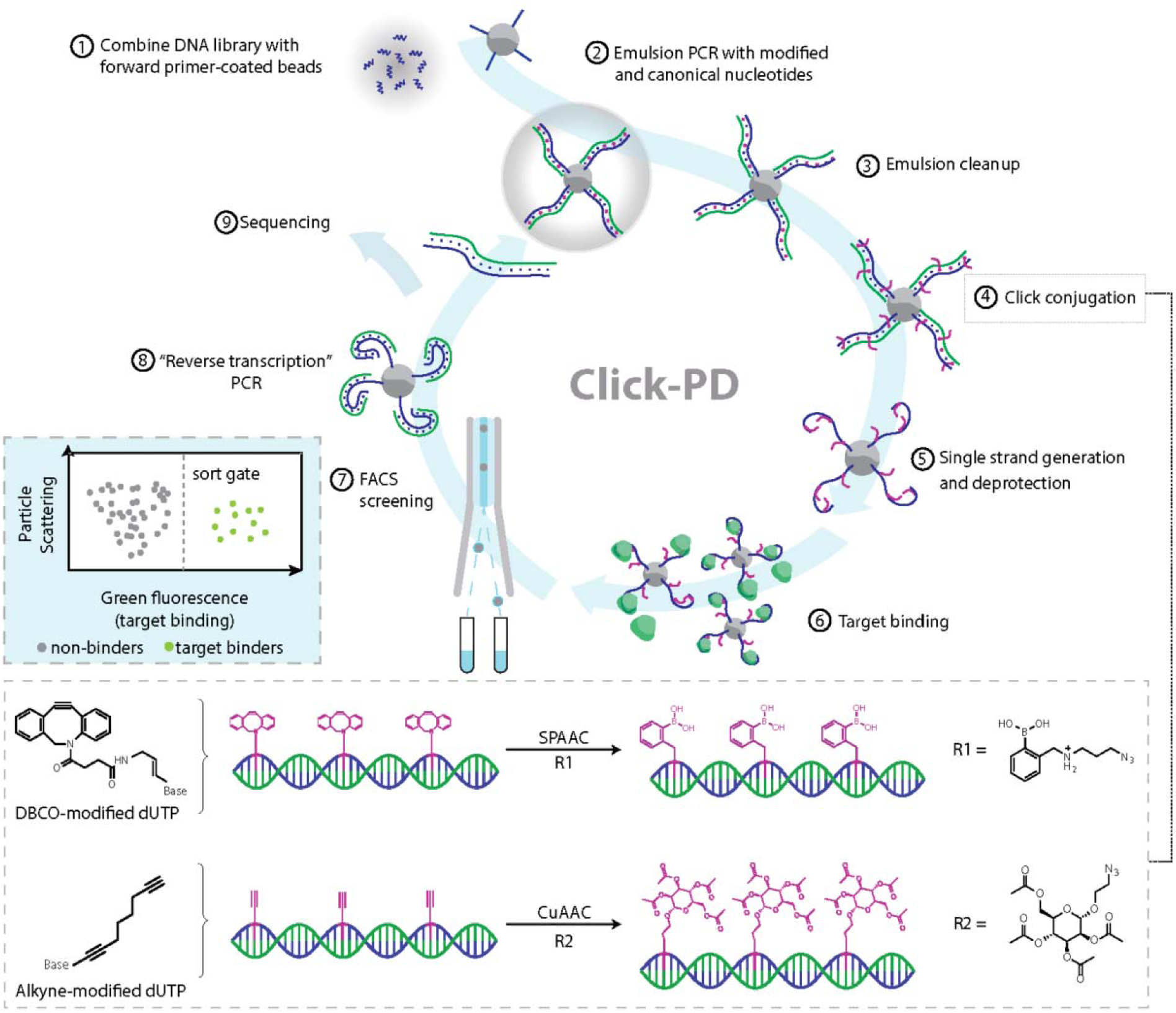
Click-PD strategy for the synthesis and screening of base-modified aptamers. After combining the initial DNA library with forward primer-coated magnetic beads (step 1), we perform emulsion PCR (step 2) to produce monoclonal aptamer particles in which dT is substituted with alkyne modified-dUTP. We then break the emulsions (step 3) and use either strain-promoted alkyne-azide cycloaddition (SPAAC) or copper(I)-catalyzed alkyne-azide cycloaddition (CuAAC) (step 4; bottom box) to conjugate azide-modified functional groups to the alkyne group on the modified uracil nucleotides. R1= azido-phenylboronic acid, R2 = 2-azidoethyl 2,3,4,6-tetra-O-acetyl-α-d-mannopyranoside. These are converted to single-stranded aptamers (step 5) containing modified deoxyuridine and combined with fluorescently-labeled target molecules (step 6). FACS screening allows us to isolate molecules that exhibit high-affinity target binding (step 7, left inset box). The selected base-modified aptamers are then converted back to natural DNA via a reverse transcription-like PCR reaction (step 8) and subjected to sequencing (step 9) or further screening.

After breaking the emulsion and removing the PCR reagents, the particles are isolated (step 3) and conjugated with an azido-labeled functional group through a SPAAC or CuAAC reaction (step 4). Nucleobase modification for the epinephrine selection was performed using SPAAC because Cu(I) is incompatible with the boronic acid modification that was used in the selection, whereas ConA selection was performed with CuAAC. The modified, double-stranded PCR products are subsequently treated with NaOH to remove the antisense strand, producing particle-displayed aptamers that incorporate chemically-modified nucleotides (step 5). We note that conjugation is performed while the products are still double-stranded. We opted for this approach because the alkyne side-chain at the 5-position of uracil adopts an outward-pointing conformation in the major groove of the double helix,^26^ which prevents steric hindrance caused by single-stranded nucleic acid folding and thus allows for more efficient and uniform modification.

We then incubate the resulting library of aptamer particles with fluorescently-labeled target (step 6) and use FACS to sort individual base-modified aptamer particles (step 7) at a throughput of ∼10^7^ particles/hour. Finally, we perform a ‘reverse transcription’ PCR reaction — again, using a commercially-available polymerase—to convert the selected base-modified aptamers back to natural DNA (step 8), with the enriched pool used for either a new round of screening or sequencing (step 9). We performed extensive testing to optimize reaction conditions and reagent selection to ensure the efficiency and sequence fidelity of the various key steps of the click-PD procedure (*i*.*e*. modified-dUTP incorporation, click modification, and reverse transcription), and have detailed these optimization experiments in the Supplementary Methods section (**Fig. S1-7**).

### Click PD generates a novel epinephrine aptamer

As our first target, we performed a click-PD screen for the small-molecule epinephrine (also known as adrenaline), for which there are no published aptamers. Aptamers for small molecules generally exhibit modest affinity, with *K*_*d*_ in the 10–100 μM range,^28^ and we aimed to generate base-modified aptamers with higher affinities for epinephrine. To this end, we chose boronic acid as our modification, since this functional group forms a reversible covalent bond with a diol group^29–32^ found in epinephrine.

We first performed a round of conventional SELEX with bead-immobilized epinephrine to pre-enrich the pool prior to the FACS screen. We also performed negative selection to eliminate sequences that bind to the particles or to the fluorescein isothiocyanate (FITC) dye used to label the epinephrine. After the negative selection, we PCR amplified the pool onto beads with DBCO-dUTP using emulsion PCR. Once our pre-enriched pool of boronic acid-modified aptamer particles was prepared, we performed four rounds (R1–4) of click-PD with FITC-labeled epinephrine. We measured binding of the aptamer particles to the target over a range of concentrations to identify the optimal target concentration for FACS. For the first round, we used 10 μM FITC-epinephrine because it resulted in 0.1% of the aptamer particles binding to the target—a level of target binding that has previously been shown to give the maximum theoretical enrichment between rounds.^16^ We observed that the binding fraction increased in successive rounds (**Fig. 2A**), and found that 1.25% of input R4 aptamer particles bound to epinephrine at 10 µM.

**Figure 2.**
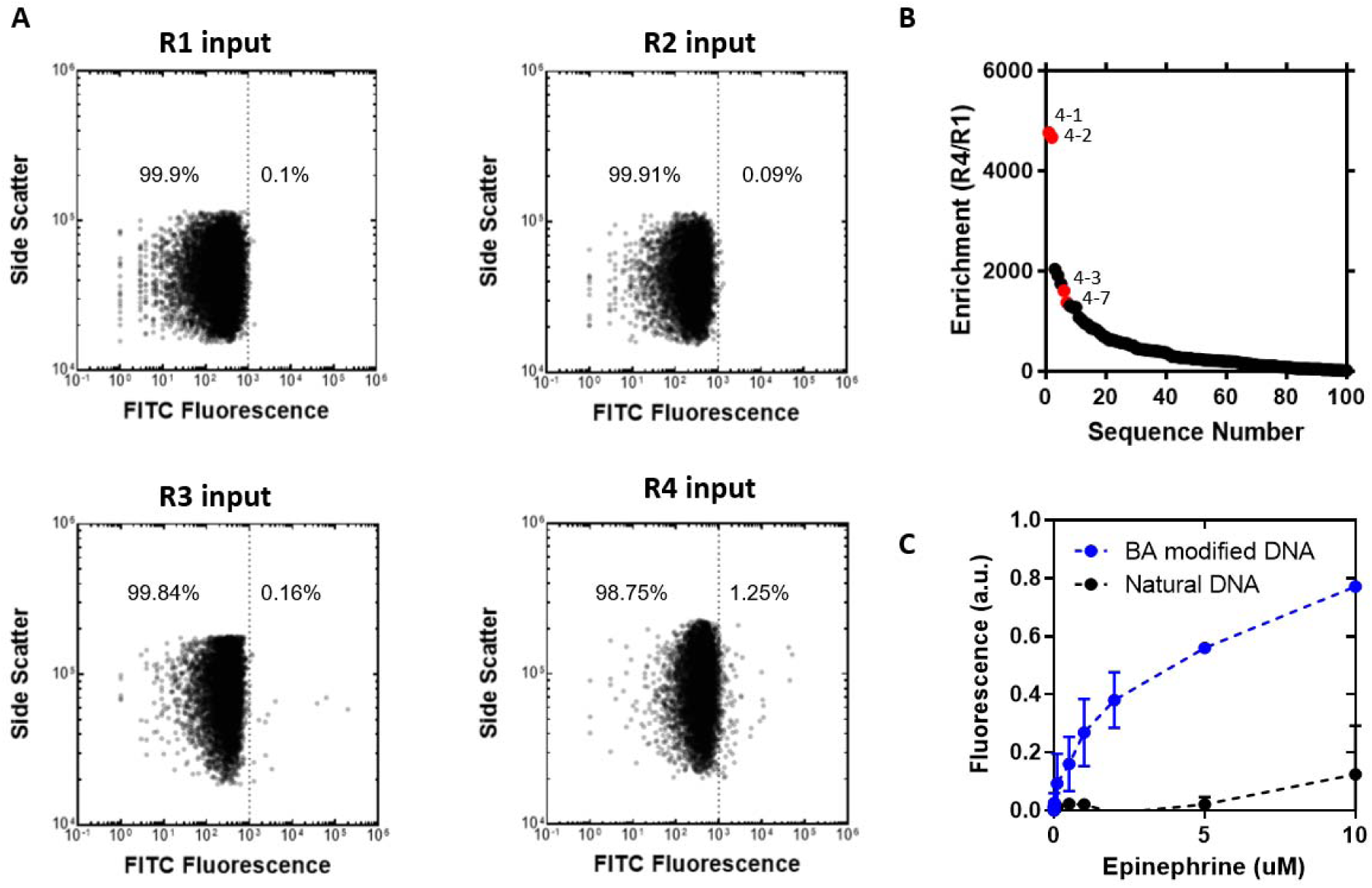
Click-PD screening for a boronic acid-modified aptamer for epinephrine. **A**) Flow cytometry analysis of aptamer particles during click-PD selection for 10 μM epinephrine over four rounds of screening. The dotted line indicates a cut-off of 1,000 a.u. FITC fluorescence; numbers indicate the percentage of particles with fluorescence above this threshold. A population of particles that display high FITC fluorescence becomes apparent in the input from R4, indicating enrichment of aptamer sequences that bind epinephrine. B) The 100 most common sequences in R4, plotted against their enrichment from R1 to R4. Sequences highlighted in red were selected for further testing. C) Fluorescent bead-based binding assay of our top epinephrine aptamer candidate, 4-1. We calculated a *K*_*d*_ of 1.1 μM using GraphPad. Without the boronic acid modification, the natural DNA shows no binding to epinephrine.

We subsequently used high-throughput sequencing with an Illumina MiSeq to identify sequences that were enriched over the course of four rounds of click-PD (See Methods). The FASTQ sequencing data was preprocessed using Galaxy NGS tools^33^ and analyzed using FASTAptamer^34^ to count and rank the unique sequences in each pool (see Supplementary Methods). As screening progressed, the population of unique sequences in the pools decreased from 95% after R1 to just 3% in the final pool. This confirms that our aptamer pools were converging towards lower diversity, with increased representation of a smaller number of aptamer sequences with affinity for epinephrine. We next calculated the enrichment factor for each aptamer sequence based on the ratio of how frequently that sequence occurred in the R4 pool versus the R1 pool, and this revealed candidates that showed up to 4,762-fold enrichment between R1 and R4 (**Figure 22B**). We also identified two major aptamer families based on sequence relationship (**Fig. S10**). Many of the top sequences fell into one of these families, differing only by single or double point-mutations. The top sequence represented 56% of the final pool. We selected two sequences from each family (**Table S2**) that had undergone >100-fold enrichment for further testing, synthesizing particles displaying each of these sequences and measuring their fluorescence intensity after incubating with 1 nM to 10 μM epinephrine (**Fig. S11**). Of these four, sequence **4-1** was selected for further characterization based on its strong binding to epinephrine.

We measured the affinity of **4-1** using a bead-based fluorescent assay. Monoclonal aptamer beads were incubated with increasing concentrations of fluorescently-labeled target and analyzed using flow cytometry, after which we normalized the mean fluorescence and generated a binding curve. We measured a K_d_ of 1.1 μM for **4-1** (**Fig. 2C**). The boronic acid modification plays an essential role in the aptamer’s target affinity, and we observed no binding when we synthesized an identical aptamer sequence composed of natural DNA. This result highlights the clear value in being able to rapidly generate and screen libraries of base-modified aptamers for challenging targets.

### Two-color click-PD generates Con A aptamers with excellent specificity

Click-PD also offers the capability to perform screening for affinity and specificity in parallel in a single screening experiment. We have previously demonstrated that particle display offers such capabilities for natural DNA aptamers by exploiting multi-color FACS sorting,^17^ and have likewise adapted this approach for use with click-PD. To demonstrate these capabilities, we chose the lectin Con A as a target because lectins pose a considerable challenge for the generation of highly specific affinity reagents. Due to the high degree of structural homology among lectins, selections tends to result in molecules with reasonable affinity but poor specificity.^35,36^ In order to achieve selection of a high-specificity aptamer with click-PD, we labeled our target, Con A, with AlexaFluor 647 and labeled a second, non-target competitor lectin, *Pisum sativum* agglutinin (PSA), with FITC. PSA is another mannose-binding lectin with considerable structural homology to Con A.^35,37^ The use of these two differentially-labeled molecules allows us to measure both on- and off-target binding simultaneously using FACS, and isolate only those aptamers that exhibit strong and selective binding for the intended target.

Since Con A preferentially binds to mannose, we chose to conjugate our nucleic acid library with a modified glycan group that mimics mannose, 2-azidoethyl 2,3,4,6-tetra-O-acetyl-α-D-mannopyranoside. Importantly, this did not require any alterations to our process, and our two-color click-PD screen employed the same library design described above for epinephrine, making use of C8-alkyne-dUTP as a handle for modification via CuAAC (**Fig. 3A**). However, we anticipated that the extensive flexible carbohydrate modifications on the nucleic acid backbone could increase the entropic penalty for forming a stable protein-aptamer complex, particularly when exposed to solvent. To mitigate this potential problem, we introduced a second nucleotide modification that could improve the binding stability. Specifically, we replaced deoxycytidine (dC) with 5-formyl-deoxycytidine, which bears an electrophilic aldehyde group that can confer potential interactions with nucleophilic groups, both intramolecularly and on the target molecule. This strategy gave rise to a large and diverse collection of three-dimensional structures that display the modification of interest in a wide range of positions and demonstrates that our approach can be used to introduce more than one distinct modification into an aptamer library.

**Figure 3.**
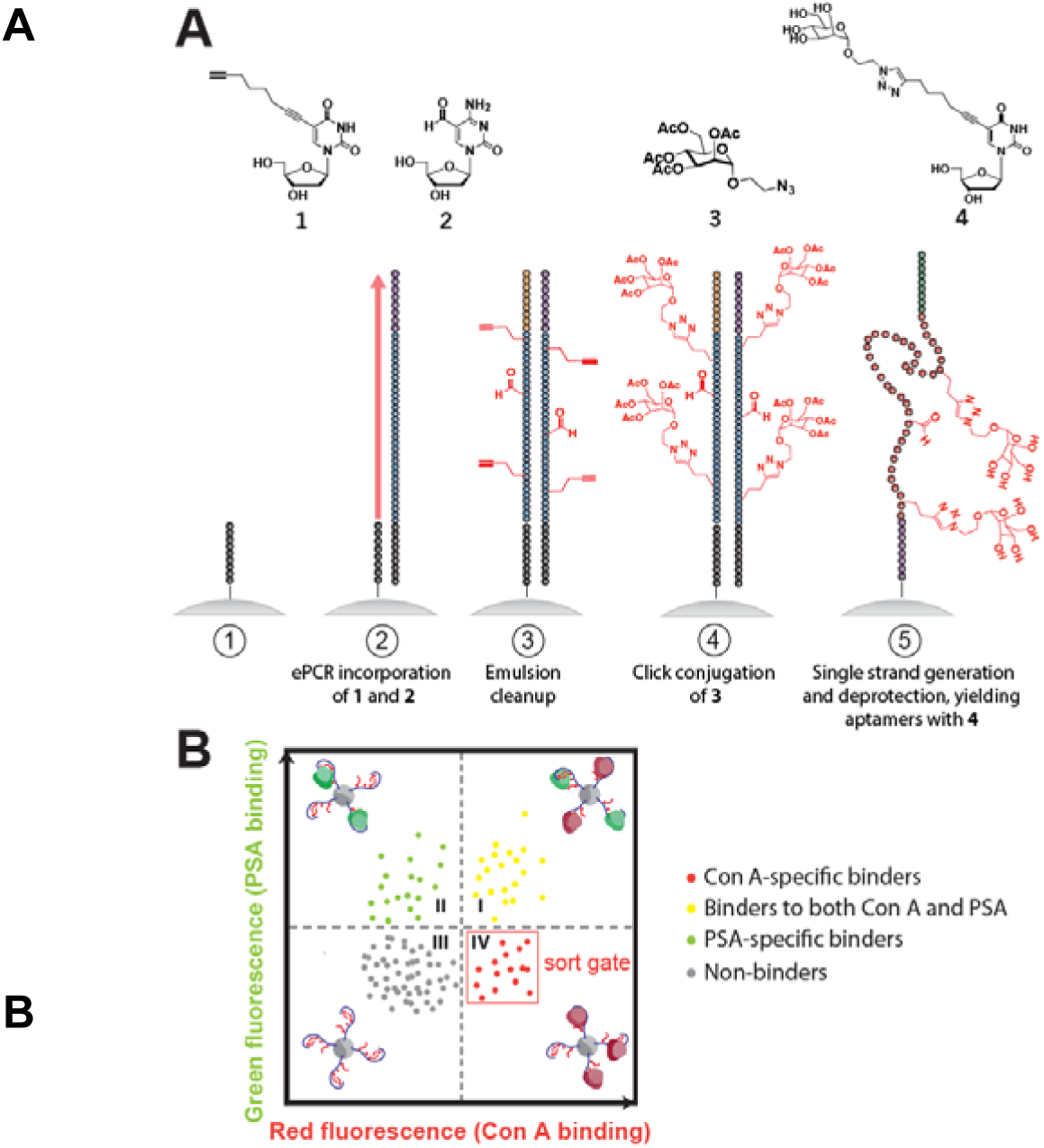
Click-PD strategy for two-color screening of lectin-specific base-modified aptamers. (**A**) Structures of modified nucleotides and chemical adducts (top), and illustration of the base-modified aptamer synthesis process (bottom). Using FP-coated magnetic beads (step 1), we performed emulsion PCR (step 2) to create monoclonal aptamer particles with C8-alkyne-dUTP and 5-formyl-deoxycytidine replacing dT and dC, respectively. After breaking the emulsions (step 3), we used a CuAAC reaction (step 4) to conjugate an azide-modified mannose derivative to alkyne-modified deoxyuridine nucleotides. Single strand generation and deprotection are then performed (step 5), yielding aptamers containing carbohydrate-modified deoxyuridine and aldehyde-modified dC. (**B**) Illustration of FACS plot with two-color screening for affinity and specificity in parallel. The aptamer particles are combined with both target and non-target lectins, each labeled with a distinct fluorophore. FACS screening allows us to exclusively isolate those molecules that exhibit strong and specific target binding (lower-right quadrant).

After incubating our aptamer particles with both Con A and PSA, we used FACS to sort individual particles that simultaneously exhibit high Alexa 647 fluorescence (and thus high Con A affinity) and weak FITC fluorescence (and thus low PSA affinity) (**Fig. 3B**). We started with ∼10^8^ base-modified aptamer-displaying particles, and a fraction of this starting population already had strong affinity for Con A at a concentration of 1 nM, even without any pre-enrichment (**Fig. 4A**). However, most of these molecules lacked specificity, as shown by their notable binding to PSA. This lack of specificity was expected, given that both lectins bind strongly to mannose.

**Figure 4.**
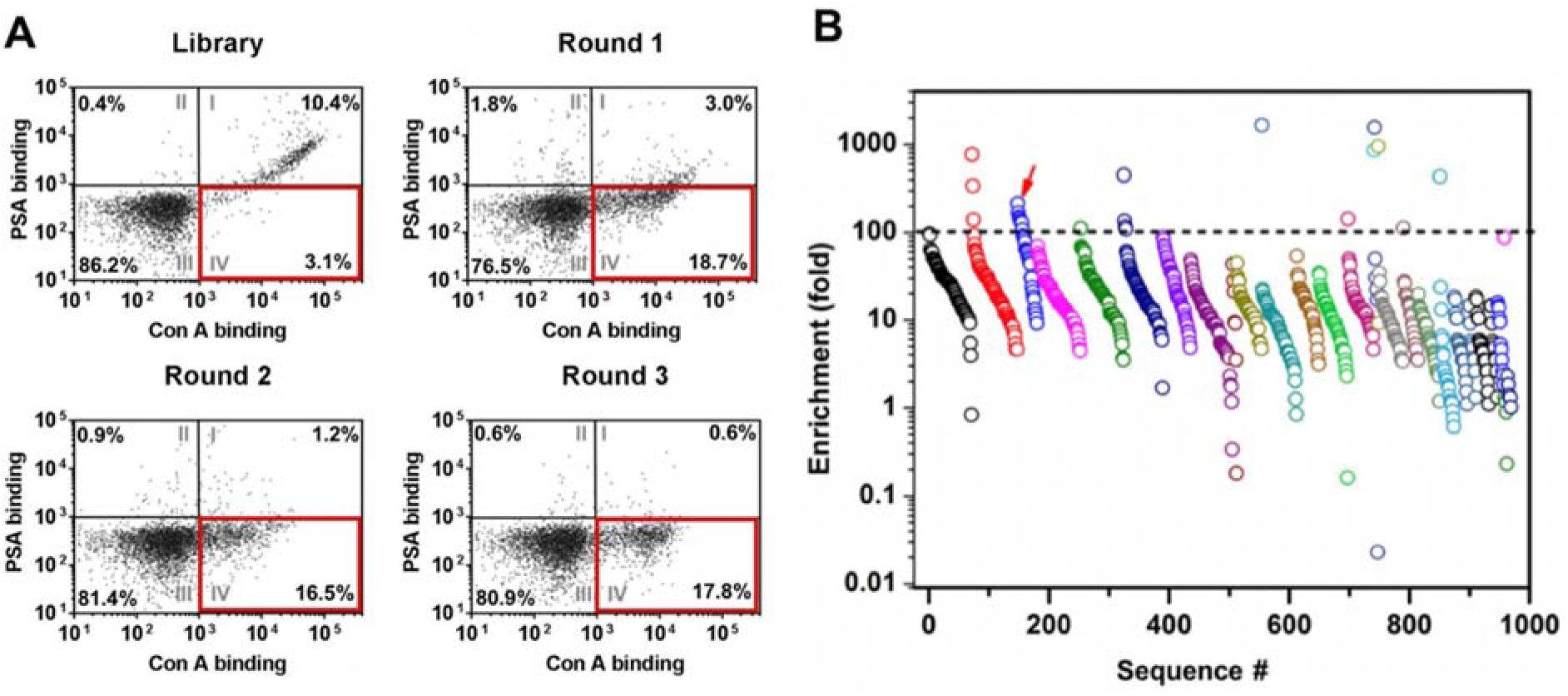
Two-color click-PD screening generates base-modified aptamers with high affinity and specificity for Con A. (**A**) FACS plots of base-modified aptamer-displaying particles from the starting library and the aptamer pools from R1–3, where [Con A] = 1 nM and [PSA] = 250 nM. Percentages represent the subpopulation of particles in each quadrant. Quadrant IV (outlined in red) represents aptamers with high Con A and low PSA affinity, which were collected in each round. (**B**) High-throughput sequencing shows several highly-enriched clusters of closely related sequences in the R3 pool. Each circle represents one enriched sequence, with colors indicating related clusters. The dotted line depicts our threshold for the most highly-enriched sequences (>100-fold). Aptamer **ConA-3-1** (red arrow) was selected for further characterization.

We performed three rounds (R1–R3) of screening, collecting only particles that exhibited strong Con A binding without binding PSA (**Fig. 4A**). We observed a clear increase in the specificity of the selected particles from round to round, and by the end of R3, 17.8% of the population bound strongly to 1 nM Con A without binding to PSA, even in the presence of a 250-fold higher concentration of the competitor.

We then performed high-throughput sequencing of the R1–3 pools to identify sequences that had become highly enriched during the click-PD process. After filtering out low-quality sequences (where >10% of bases had a quality score ≤20) using Galaxy NGS tools^33^ (see Supplementary Methods), we obtained 182,499 unique sequences (684,179 reads) in the R1 pool, 150,680 unique sequences (643,462 reads) in the R2 pool, and 2,867 unique sequences (470,426 reads) in the R3 pool. We identified 132 sequence clusters, which we defined as groups of closely-related sequences that differ from one another by two or fewer mutations,^34^ in the R3 pool. The degree of enrichment from R1 to R3 varied for the sequences within each cluster, with some of the most highly-enriched clusters containing sequences that had undergone 100-fold to >1000-fold enrichment (**Fig. 4B**). We selected 14 sequences exhibiting >100-fold enrichment for further testing, synthesizing particles displaying each of these sequences and measuring their fluorescence intensity after incubating with 1 nM Con A (**Table S3**). Sequence **ConA-3-1** (**Table S1**) was selected for further characterization due to its strong binding to Con A and the fact that it belonged to a highly enriched (>2,000-fold) sequence cluster (**Fig. S12**).

### ConA-3-1 target binding is modification-dependent

We subsequently demonstrated the excellent affinity and specificity of **ConA-3-1** for its target lectin. We incubated particles displaying **ConA-3-1** with different concentrations of fluorescently-labeled Con A and PSA, and then measured the fluorescence intensity of the particles using FACS. We established a binding isotherm by plotting the percentage of target-bound particles over the total population at each lectin concentration. This revealed strong affinity for Con A (*K*_*d*_ = 20 nM) and much weaker affinity for PSA (*K*_*d*_ > 1 µM), clearly demonstrating the excellent specificity of this molecule (**Fig. 5A**).

**Figure 5.**
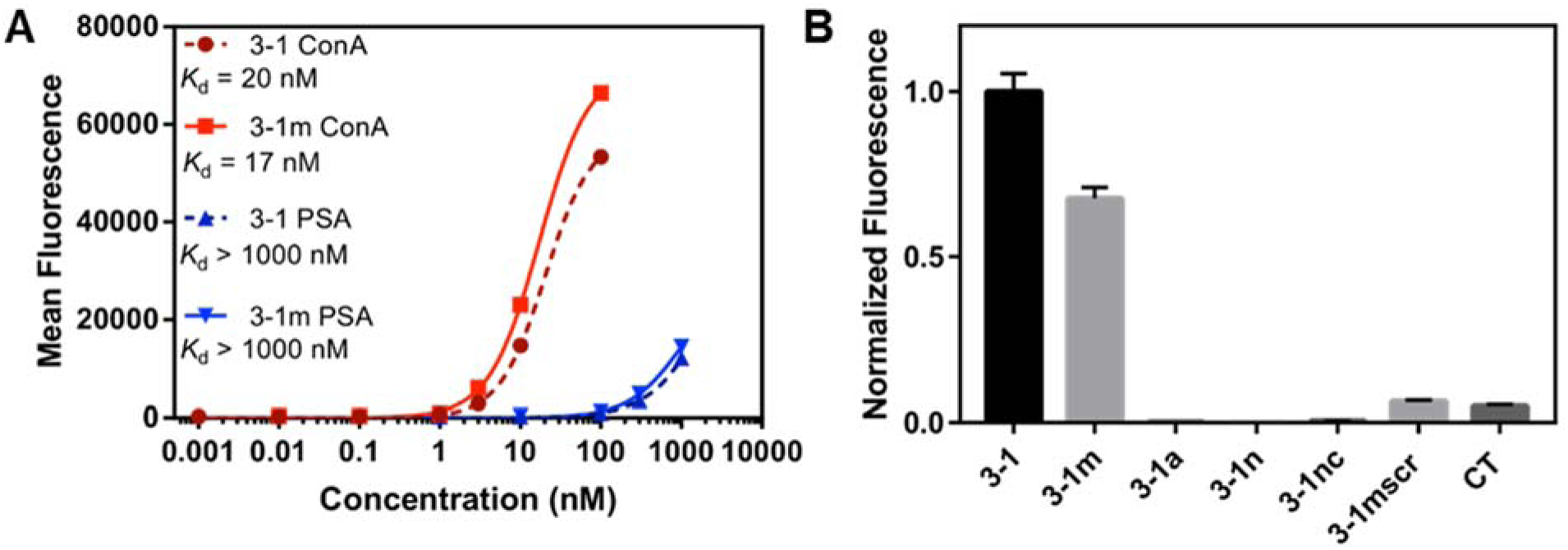
Contribution of modifications to Con A aptamer affinity. (**A**) Binding curves of **ConA-3-1** and **ConA-3-1m** to Con A and PSA from particle-based fluorescent measurements. (**B**) Binding activity for various **ConA-3-1** derivatives in the presence of 10 nM Con A. Fluorescence intensities were normalized first to particle coating, then to the relative signal of **ConA-3-1**.

**ConA-3-1** contains multiple modifications, and we sought to determine the extent to which these modifications contribute to its strong and specific interaction with Con A. We synthesized particles displaying various mutant sequences based on **ConA-3-1** with different modification profiles (**Table S1**). **ConA-3-1a** no longer contained uridine or mannose modifications, but still displayed aldehyde groups. On the other hand, **ConA-3-1m** lacked aldehyde groups but still had mannose modifications. We also prepared a construct composed entirely of canonical bases (**ConA-3-1n**), and a version of **ConA-3-1** that was not subjected to subsequent click conjugation (**ConA-3-1nc**). Finally, to confirm that the affinity of **ConA-3-1** is sequence-specific, we prepared a ‘CT-only’ sequence (**CT**) that was the same length as **ConA-3-1** but only contained dC and mannose-modified C8-alkyne-dUTP, where the latter were present in a number equal to that of **ConA-3-1**, and a sequence with the same nucleotide composition as **ConA-3-1m** but in a scrambled order (**ConA-3-1mscr**).

**ConA-3-1a, ConA-3-1n**, and **ConA-3-1nc** showed essentially no binding to 10 nM Con A (**Fig. 5B**), indicating that Con A binding was mannose-dependent. Both **CT** and **ConA-3-1mscr** showed only low levels of binding to 10 nM Con A, which is most likely attributable to the presence of mannose functional groups. Notably, **ConA-3-1m** showed only slightly lower levels of binding to 10 nM Con A than **ConA-3-1**, despite the absence of aldehyde modifications (**Fig. 5B**). This unexpected finding prompted us to further investigate **ConA-3-1m**’s binding profile. We determined that the affinity of **ConA-3-1m** for Con A is in fact slightly superior to that of **ConA-3-1** (*K*_*d*_ = 17 nM), and that the absence of aldehyde-modified dC did not affect **ConA-3-1m**’s specificity for PSA (*K*_*d*_ > 1 µM) (**Fig. 5A**). This indicates that the aldehyde functional groups do not contribute meaningfully to **ConA-3-1**’s affinity or specificity, and that only the mannose modifications are absolutely required for Con A binding.

We further validated the binding characteristics of **ConA-3-1** and **ConA-3-1m** by using an alternative measurement method, bio-layer interferometry (BLI).^38^ This allowed us to confirm that these binding results are independent of the particles on which the aptamers are immobilized, and to measure association rate (*k*_*on*_) and dissociation rate (*k*_*off*_) constants. Solution-phase base-modified aptamers were prepared using conventional PCR instead of emulsion PCR, with biotinylated FP instead of particle-conjugated FP, and with ESI-MS confirmation after click conjugation with the mannose group (**Fig. S13**). We immobilized biotinylated **ConA-3-1** and **ConA-3-1m** onto the streptavidin-coated surface of the biosensor and incubated with Con A at various concentrations, followed by dissociation in blank buffer (**Fig. S14**). For **ConA-3-1m**, we globally fitted the resulting response curves for each concentration to generate rate constants of *k*_*on*_ = 7.1 ± 0.3 × 10^4^ M^−1^ s, and *k*_*off*_ = 2.3 ± 0.02 × 10^−4^ s^−1^, corresponding to a *K*_*d*_ of 3.2 ± 0.2 nM. Notably, *k*_*off*_ for both **ConA-3-1** and **ConA-3-1m** when bound to Con A was comparable to or lower than that of many antibody-antigen interactions.^39,40^ We also fitted the maximum response measurements from each concentration to a cooperative binding model, yielding a *K*_*d*_ of 5.3 ± 0.7 nM for **ConA-3-1m**. These affinity values are in reasonable agreement with the measurement from our particle-based binding assay. In comparison, the *K*_*d*_ of **ConA-3-1** for Con A is 5.8 ± 0.8 nM by BLI (**Fig. S14**), confirming that the substitution of dC with 5-formyl-deoxycytidine does not enhance lectin binding, and indeed slightly reduces affinity in the BLI assay.

To further determine the extent to which each mannose side-chain contributes to **ConA-3-1m**’s interaction with Con A, we generated particles displaying mutants of **ConA-3-1m** in which either individual occurrences or pairs of mannose-conjugated deoxyuridine within the sequence (excluding the primer region) were substituted with dA, and screened their affinity for Con A (**Fig. S15**). We determined that essentially all of the mannose groups, with the exception of those at nucleotide positions 45 and 46, contribute to binding, and that the loss of even one mannose side-chain in the sequence significantly reduced the affinity of the mutant.

### ConA-3-1m demonstrates excellent specificity

Having shown **ConA-3-1m**’s strong specificity for Con A versus PSA, we subsequently demonstrated its ability to discriminate against a wide variety of other closely-related lectins that also preferentially bind mannose. Plant-derived mannose-binding lectins such as *Lens culinaris* agglutinin (LcH), *Narcissus pseudonarcissus* lectin (NPA), and *Vicia faba* agglutinin (VFA) all belong to the same carbohydrate specificity group as Con A and PSA and share high structural homology,^41^ and are therefore good models for testing specificity. Critically, **ConA-3-1m** showed virtually no binding to LcH, NPA, or VFA at 10 nM (**Fig. S16**). Even at a 100-fold higher concentration (1 μM), **ConA-3-1m** showed little binding to LcH and NPA, and only modest binding to VFA. This low level of binding to VFA at high concentrations can be attributed to the especially high degree of homology between Con A and VFA.^41^

Next, we expanded our analysis of **ConA-3-1m** to a group of 70 structurally related and unrelated lectins using a lectin array (**Fig. 6A**) that included lectins belonging to different specificity groups with varying degrees of homology to Con A (**Table S4**). This assay further confirmed the remarkable specificity of **ConA-3-1m**. Across a broad range of concentrations from 0.04–400 nM, Con A generated the strongest signal among all the lectins (**Fig 6B**), and most produced no measurable signal.

**Figure 6.**
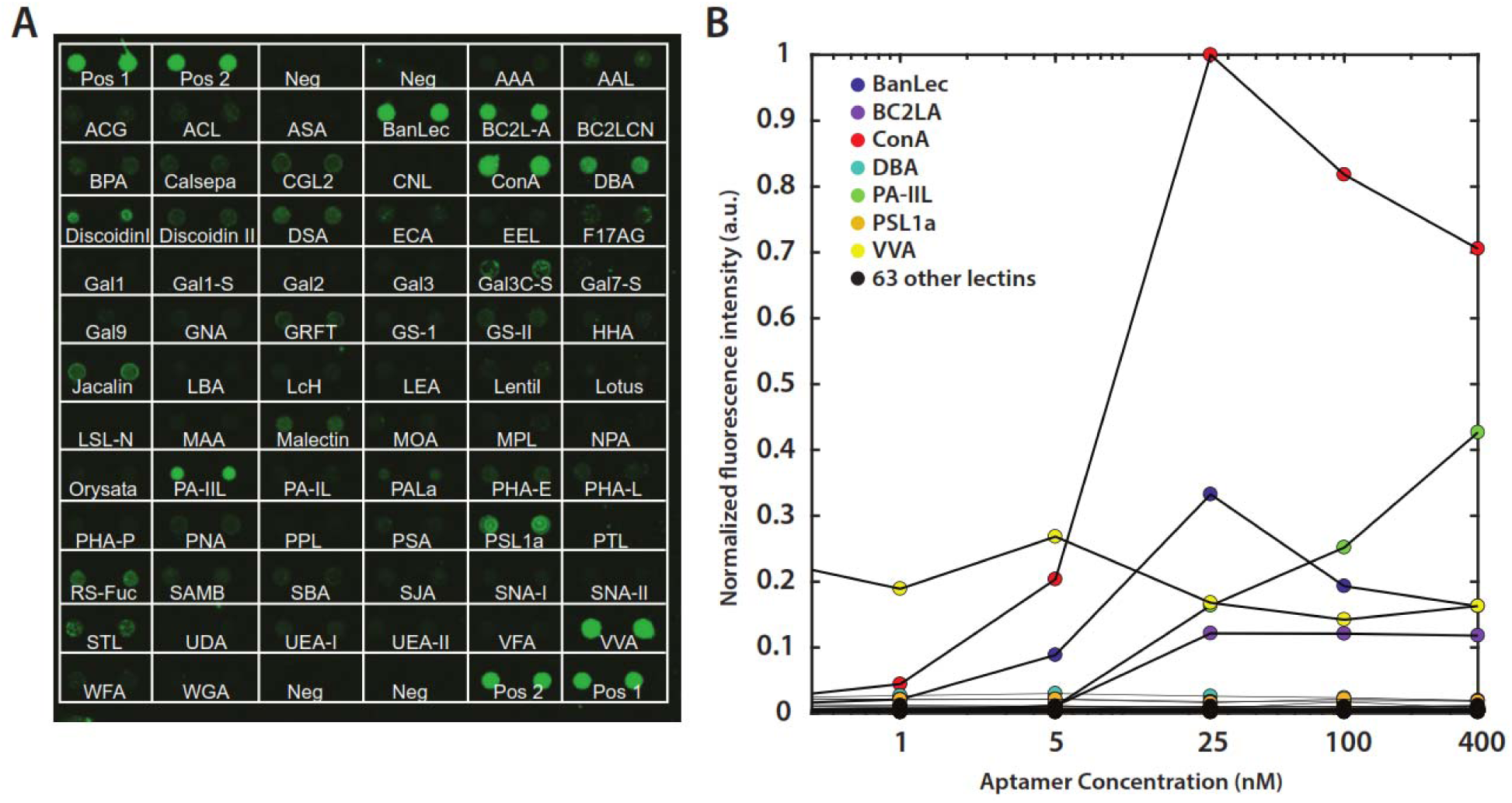
**(A)** The strong specificity of **ConA-3-1m** is clearly apparent on an array of 70 lectins incubated with 100 nM aptamer. Each lectin is spotted in duplicate. The short names of the lectins are written under the spots (detailed information in **Table S4)**; pos and neg denote positive and negative controls, respectively. (**B**) Quantitation of binding of **ConA-3-1m** to the array at lectin concentrations ranging from 40 pM to 400 nM. Data are normalized to the maximum fluorescence intensity of Con A. Positive and negative control array data were not included in the plot. The slightly lower fluorescence signal seen with Con A at the two highest concentrations was attributed to the self-quenching effect of the fluorophores at high local concentrations on the array surface.

**ConA-3-1m** showed binding activity to six other lectins on the array: BanLec, BCL2-A, DBA, PA-IIL, PSL1a and VVA **(Fig 6A & Fig S17A)**. BanLec^42^, BC2L-A^43^, PA-IIL^44^ all have been reported to exhibit affinity for mannose. For DBA, PSL1a, and VVA, the signal did not change meaningfully in response to increasing concentration of aptamer, indicating that these likely represent false-positive binding events (**Fig 6B**). We repeated this assay with a previously published ConA aptamer, ^45^ and saw barely any binding to Con A (**Fig. S17B)** and no evidence for meaningful target specificity, as this aptamer bound to all the other lectins with similar affinity (**Fig. S17C**).

### ConA-3-1m shows potent inhibition of erythrocyte agglutination

Given the strong affinity and remarkable specificity of our base-modified aptamer for Con A, we hypothesized that it might act as a highly effective inhibitor of Con A’s biological activity. Con A induces “clumping” of human erythrocytes in a process known as hemagglutination,^46^ a standard assay for quantifying activity of this lectin. As a baseline, we established that complete hemagglutination occurs at 150 nM Con A, based on visual observation of the deposition of erythrocytes in a 96-well plate. This was confirmed by monitoring absorbance of the cell suspension at 655 nm, which correlates to the size of the agglutinated clump.^47^ We then tested the extent to which **ConA-3-1m** can inhibit this process by incubating various concentrations of **ConA-3-1m** with 150 nM Con A for 30 min at room temperature before adding erythrocytes at 1% hematocrit. We observed concentration-dependent inhibition of Con A-induced hemagglutination, with complete inhibition at 150 nM and a half-maximal inhibitory concentration (IC50) of 95 nM (**Fig. 7A, B**). The observation that complete inhibition occurs when both Con A and **ConA-3-1m** are at the same concentration (150 nM) indicates a stoichiometric relationship, confirming the strong and stable interaction between the binding partners.

**Figure 7.**
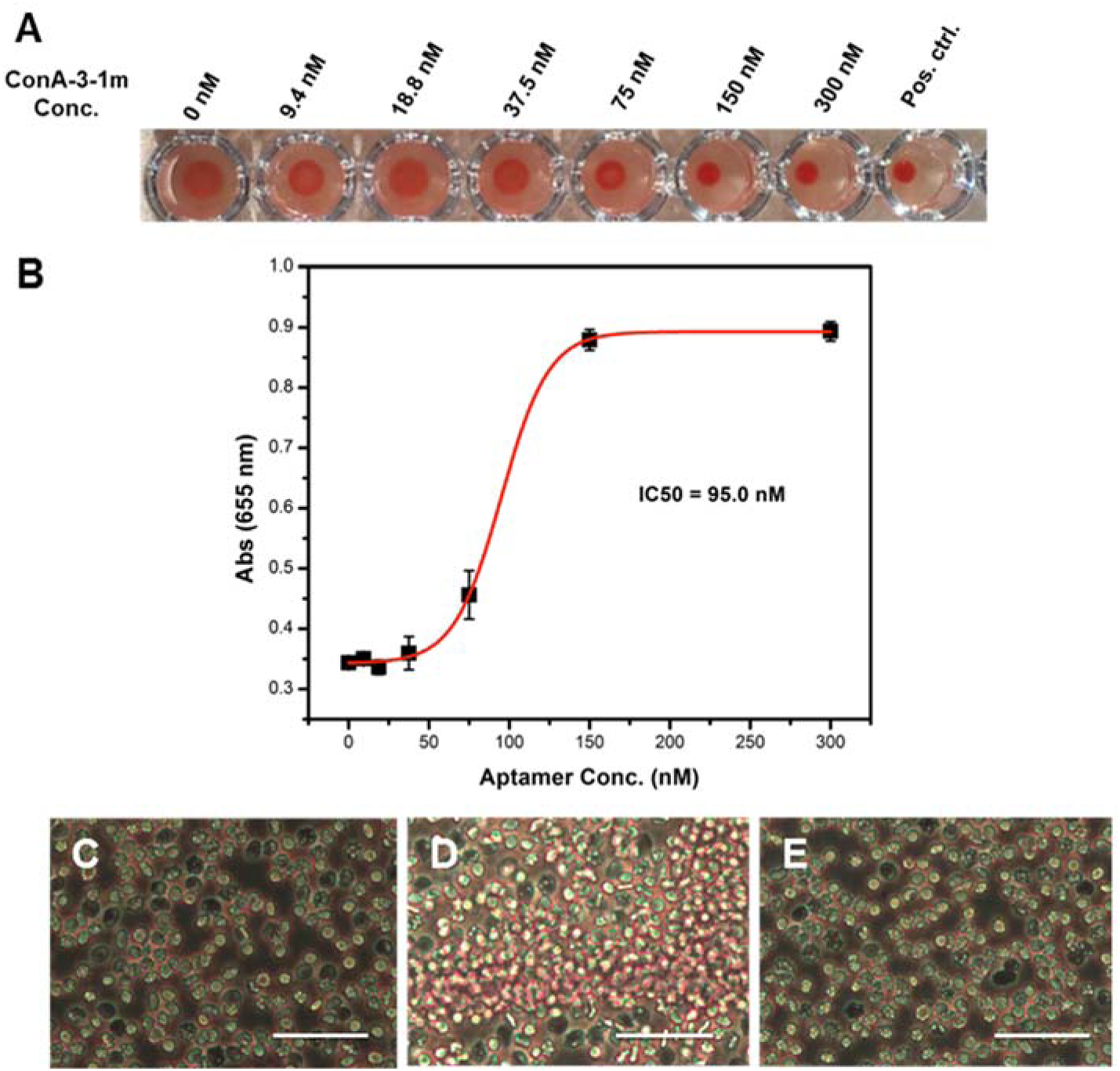
ConA-3-1m is a potent inhibitor of Con A-induced hemagglutination. (**A**) We incubated various concentrations of **ConA-3-1m** with a human erythrocyte suspension containing 150 nM Con A, a concentration known to induce complete hemagglutination. The deposition of erythrocytes onto the bottom of the wells indicates inhibition of Con A activity. The positive control well contains only human erythrocytes, with no Con A. (**B**) Inhibition of hemagglutination, as measured by increased absorbance at 655 nm. We observed that **ConA-3-1m** inhibited 150 nM Con A with an IC50 of 95.0 nM. Error bars were derived from four replicates. (**C**–**E**) 40X microscopic images of normal human erythrocytes (**C**) and human erythrocytes incubated with (**D**) 0.65 µM Con A or (**E**) 0.65 µM Con A with 0.8 µM **ConA-3-1m**. Scale bars = 40 µm.

We also microscopically monitored inhibition of Con A-induced hemagglutination by **ConA-3-1m** (**Fig. 7C–E**); the erythrocyte clumps that formed upon the addition of Con A were absent when we incubated Con A with **ConA-3-1m** beforehand. Notably, **ConA-3-1m** inhibits Con A-induced hemagglutination with ∼10^7^-fold greater potency than methyl α-D-glucopyranoside, a commonly used inhibitor that achieves maximal effect at 50 mM.^48^ Furthermore, **ConA-3-1m** is about three-fold more potent than the best-known inhibitor described to date for Con A, a mannose glycopolymer reported by Kiessling *et al*., which achieves complete inhibition at 500 nM. This is particularly striking given that **ConA-3-1m** contains 120-fold fewer mannose side chains (14 units) compared with the mannose glycopolymer (∼1,700 units), suggesting that its carbohydrate presentation more closely aligns with the active sites of this lectin.^22^

## CONCLUSION

The use of non-natural, modified nucleotide analogues can greatly expand the chemical and functional space available to aptamers, but efforts to isolate such molecules have previously been impeded by the need to engineer polymerase enzymes that can efficiently process these various modified nucleotides. As a solution to this problem, we have coupled a simple and robust click chemistry-based DNA modification strategy with our particle display screening platform to develop click-PD, which enables the efficient generation and high-throughput screening of diverse base-modified DNA aptamers that can incorporate a wide range of functional groups. As proof of principle, we generated a novel boronic acid-modified aptamer with ∼1 μM affinity for epinephrine —the first aptamer described to date for this target. We subsequently performed a two-color click-PD screen to generate a mannose-modified aptamer with low nanomolar affinity for the lectin ConA. Our **ConA-3-1m** aptamer exhibited exceptional specificity, with minimal binding to other structurally similar lectins, and also showed remarkable biological activity—to the best of our knowledge, it is the most potent inhibitor of Con A-mediated hemagglutination reported to date.^22^ The strong affinity and specificity of each of these aptamers were fundamentally dependent upon the presence of the incorporated chemical modifications, and we show that the removal of boronic acid or mannose groups essentially eliminated aptamer binding to epinephrine and Con A, respectively.

This method should be readily accessible to the broader research community, as it is minimally demanding in terms of specialized reagents or instrumentation. The amplification and reverse-transcription steps of click-PD are both compatible with commercially-available polymerases, and the alkyne-modified nucleotide employed here can readily be covalently coupled to any number of azide-tagged functional groups via an efficient click chemistry reaction. In terms of equipment, click-PD requires only a simple, commercially-available emulsifier apparatus and a FACS machine—instrumentation that is now available at many research institutions. As such, we believe that click-PD offers a powerful platform for efficiently generating custom reagents for a wide range of biological and biomedical applications.

## ASSOCIATED CONTENT

Supporting Information

Supplementary Methods and Supplementary Results are presented.

## ACKNOWLEDGMENT

The authors gratefully thank Dr. Gurpreet Sekhon for his help with the graphics. High-throughput sequencing was performed by the Stanford Functional Genomics Facility. We also appreciate the facility support of UCSB Biological Nanostructures Laboratory at CNSI.

