## Supplemental for "Click-PD: A Quantitative Method for Base-Modified Aptamer Discovery"

### I. Supplementary Methods

##### General methods

All DNA oligonucleotides were purchased from Integrated DNA Technologies. Primers were ordered with standard desalting. PCR templates were ordered with PAGE purification. Other than the exceptions noted below, all commercially available reagents and lab supplies were purchased from Sigma-Aldrich. 2-azidoethyl 2,3,4,6-tetra-O-acetyl-α-D-mannopyranoside and 2-azidoethyl 2,3,4,6-tetra-O-acetyl-β-D-galactopyranoside were purchased from LC Scientific. Dynabeads MyOne carboxylic acid and streptavidin C1 beads, KOD-XL DNA polymerase, MES buffer (pH 4.7), and methyl-$(PEG)_{12}$-amine were purchased from Thermo Fisher Scientific. Taq polymerase was purchased from Promega. Pwo DNA polymerase was purchased from Roche. C8-Alkyne-dUTP was purchased from Axxora. DBCO-$PEG_4$-dUTP was purchased from Jena Bioscience. 5-formyl dCTP was purchased from Trilink Technologies. Deep Vent DNA polymerase and standard dNTPs were purchased from New England Biolabs. Azido-dPEG-amine was obtained from Quanta Biodesign. Fluorescein isothiocyanate PEG NHS (FITC-PEG-NHS, $M_w$ 5000) was purchased from NANOCS. Lectin Array 70 was purchased from RayBiotech, Inc. Human erythrocytes were purchased from BioIVT. Mini-PROTEAN native and denaturing PAGE gels (10%) were purchased from Bio-Rad. Flow cytometry assays were performed using a BD Accuri C6 flow cytometer. Fluorescence-based sorting of particles was done using a BD FACSAria III.

##### Optimizing polymerase-mediated incorporation of modified pyrimidine building blocks

We performed a series of experiments to select an appropriate polymerase for base-modified aptamer synthesis. The Con A screen is particularly complex, as it requires the incorporation of two modified nucleotides and entails a more challenging monosaccharide modification. We screened several DNA polymerases to identify a candidate that allows effective replacement of dT and dC with C8-alkyne-dUTP and 5-formyl-deoxycytidine, respectively, during PCR. We used a PCR mixture containing 1X polymerase buffer, 0.2 mM dATP, 0.2 mM dGTP, 0.2 mM 5fdCTP, 0.2 mM C8-Ak-dUTP, 0.4 μM **T-FP**, 0.4 μM **T-RP**, 0.05 U/μL DNA polymerase (KOD-XL, Pwo, or Deep Vent), 20 pM PCR template **T1**, and water for a total volume of 50 μL. The cycling conditions were as follows: 96 °C for 2 min; 30 cycles of 96 °C for 15 s, 51 °C for 30 s, and 72 °C for 30 s; 72 °C for 2 min; and hold at 4 °C. We then loaded 2 μL of each PCR reaction directly onto a 10% native PAGE gel, which was run at 150 V for 30 min in 1X TBE buffer. Gels were imaged after staining with 1X GelStar Nucleic Acid Stain (Lonza) in TBE buffer. Our analysis of the final reaction (**Figure S1**) indicated that KOD-XL provides the highest yield and purity.

The same experiments was performed to determine the appropriate polymerase to incorporate DBCO-PEG4-dUTP, and our analysis of the final reaction (gel not shown) likewise indicated that KOD-XL provides the highest yield and purity.

**Optimization of CuAAC click conjugation reaction**

Next, we optimized reaction conditions for coupling mannose to C8-alkyne-dUTP via click chemistry. 10 μL of 100 μM 21-nt oligonucleotide substrate (containing three consecutive C8-alkyne-dUTP nucleotides, **Table S1**, Sequence **SI**), 1 μL 100 mM of protected sugar (2-azidoethyl 2,3,4,6-tetra-O-acetyl-β-D-galactopyranoside or 2-azidoethyl 2,3,4,6-tetra-O-acetyl-α-D-mannopyranoside); or the same amount of non-protected sugar (6-azido-6-deoxy-D-galactose or α-D-azidomannopyranoside) in DMSO (100 eq), and 14 μL 20 mM sodium phosphate buffer, pH 8 (pre-degassed by bubbling $N_2$ through) were combined in a 1.5 mL Eppendorf tube.

Click chemistry was initiated by one of the following three conditions: **(1)** addition of premixed 1 μL 20 mM $CuSO_4$, 1 μL 0.1 M tris(3-hydroxypropyltriazolylmethyl)amine (THPTA), and 20 μL water, followed by 1 μL 0.2 M sodium ascorbate. **(2)** addition of premixed 1 μL 20 mM $CuSO_4$ and 1 μL 20 mM tris[(1-benzyl-1H-1,2,3-triazol-4-yl)methyl]amine (TBTA)] in 10 μL of 4:3:1 water:DMSO:t-BuOH, followed by addition of 2 μL 20 mM tris(2-carboxyethyl)phosphine (TCEP). **(3)** addition of 10 μL premixed 1:1 Cu:TBTA (2 mM, prepared from 1 mg CuBr + 0.7 mL 10 mM TBTA in 4:3:1 water:DMSO:t-BuOH, then diluted five-fold with the same solvent).

The cap of the tube was then removed, and the de-capped tube was immediately placed in a 20 mL vial equipped with a rubber septum, followed by Ar flushing for 5 min. We incubated the sealed vial in the dark for two hours. The reaction product was purified with a Centri-Spin 10 column (Princeton Separations). 200 μL of concentrated ammonium hydroxide (18 M) was added to the purified product, and the solution was incubated at room temperature for 3 hours. 400 μL n-butanol was then added, vortex mixed, and centrifuged at 16,000 ×*g* at 4 °C for 2 min. The top organic layer was removed and discarded. The bottom aqueous layer was purified by an Oligo Clean and Concentrator spin column (Zymo Research), followed by HPLC analysis.

Reverse-phase HPLC analysis was performed on an Agilent 1100 system using a PLRP-S 4.6×150 mm 5 μm column with 300 Å packing material, with a gradient from 95% 0.1 M triethylammonium acetate (TEAA)/5% acetonitrile to 20% 0.1 M TEAA/65% acetonitrile over 30 min. We determined that the reaction conditions described in (3) above achieved quantitative yield of the fully-conjugated product for 2-azidoethyl 2,3,4,6-tetra-O-acetyl-α-D-mannopyranoside (**Figure S2**). We confirmed the successful and efficient PCR incorporation of C8-alkyne-dUTP and 5-formyl-deoxycytidine and subsequent click chemistry modification by denaturing polyacrylamide gel electrophoresis (PAGE; **Figure S3a**) and electrospray ionization mass spectrometry (ESI-MS, **Figure S3b**). ESI-MS characterization was performed by Novatia.

**Synthesis of *N*-methyliminodiacetic acid (MIDA)-protected p-azidomethyl phenylboronic acid (p-AMPBA)**

Following a modified procedure based on literature[1] (Figure S4), a 50 mL round bottom flask was charged with *p*-azidomethylphenyl boronic acid pinacol ester (0.26 g, 1 mmol), 3 mL 1 M HCl aqueous solution, polymer-bound boronic acid (3.5 g, 2.6 mmol/g, 9 mmol), and 18 mL acetonitrile. This mixture was stirred at room temperature for 36 h, with reaction progress monitored by thin layer chromatography (1:1 hexane:ethyl acetate as eluent). After the reaction was complete, the solvent was evaporated *in vacuo*, and 5 mL water was added. The crude sample was frozen by liquid nitrogen and lyophilized overnight. The residue was added into a 500 mL flask, along with MIDA (0.15 g, 1 mmol), 18 mL benzene, and 2 mL DMSO. The flask was then fitted with a Dean-Stark trap and a reflux condenser, and the mixture was refluxed with stirring for 16 h followed by concentration *in vacuo*. The resulting crude product was adsorbed onto Florisil gel from an acetonitrile solution. The resulting powder was dry-loaded on top of a silica gel column slurry-packed with ethyl acetate. The product was eluted using a gradient (2:1 ethyl acetate -> ethyl acetate:acetonitrile) to yield boronate ester as a colorless, crystalline solid (0.25 g, 87% yield over two steps). $^1$H NMR (500 MHz, $CDCl_3$): δ 7.56 (d, 2H), 7.40 (d, 2H), 4.43 (s, 2H), 4.10 (d, 2H), 2.52 (s, 3H); $^{13}$C NMR (125 MHz, $CDCl_3$): δ 168.5, 137.0, 133.0, 127.9, 117.4, 61.8, 54.2, 47.5 (**Figure S9**). HRMS (m/z) $[M+H]^+$ calculated for $C_{12}H_{13}BN_4O_4$, 288.1030, found 288.1023.

**Optimization of SPAAC click conjugation reaction**

We optimized the reaction conditions for coupling boronic acid to DBCO-PEG4-dUTP via SPAAC using sequence **T-29-DBCO** (**Table S1**). 1 µl 1 mM **T-29-DBCO** was added to 10 µl 2.78 mM p-AMPBA in 1X PBS, 100 mM NaCl and incubated overnight. Excess p-AMPBA was removed using a 3k Amicon filter and deionized water. Half of the reacted product was deprotected with 0.5M NaOH for 10 minutes at room temperature. Both deprotected and protected products were washed with deionized water. Addition of the protected boronic acid moiety and removal of the protecting group were determined by gel-shift assay (gel not shown).

**Optimization of PCR amplification, click conjugation of 2-azidoethyl 2,3,4,6-tetra-O-acetyl-α-D-mannopyranoside, single-strand generation, and acetyl deprotection**

We subsequently optimized the PCR incorporation of modified nucleotides by preparing a PCR mixture containing 1X KOD-XL DNA polymerase buffer, 0.2 mM dATP, 0.2 mM dGTP, 0.2 mM 5-formyl-deoxycytidine, 0.2 mM C8-alkyne-dUTP, 0.4 µM **T-FP**, 0.4 µM 5'-double biotinylated **T-RP-2Bio**, 0.05 U/µL KOD-XL DNA polymerase, 20 pM PCR template **T1**, and water in a total volume of 5 mL in a 96-well plate. Cycling conditions were as follows: 96 °C for 2 min; 12 cycles of 96 °C for 15 s, 51 °C for 30 s, and 75 °C for 30 s; 75 °C for 2 min; and hold at 4 °C. PCR reactions were then transferred into a 50 mL conical tube. 0.5 mL 3 M sodium acetate (pH 5.2) and 13.75 mL of 100% ethanol were added, followed by freezing at -80 °C for 30 min. The frozen stock was then centrifuged for 30 min at 21,000 × g at 4 °C to precipitate the DNA. The pellet was dissolved with 600 µL water, followed by purification using Qiagen MinElute spin columns. The PCR product was eluted with 180 µL of 10 mM Tris buffer, pH 8.0. To this DNA solution, we added 40 µL of 3 M sodium acetate (pH 5.2) and 1.2 mL of 100% ethanol, followed

[1] Gillis E. P.; Burke, M. D. *J. Am Chem. Soc.* 2008, 130, 14084-14085.

by freezing at -80 °C for 30 min. The frozen stock was then centrifuged for 30 min at 21,000 ×*g* at 4 °C to precipitate the DNA. The material was resuspended in 20 µL 1X PBS buffer.

We combined 20 µL of 100 mM 2-azidoethyl 2,3,4,6-tetra-O-acetyl-α-d-mannopyranoside in DMSO (100 eq) and 40 µL of 20 mM sodium phosphate buffer, pH 8 (pre-degassed by bubbling $N_2$ through) with 20 µL of base-modified DNA solution in a 1.5 mL Eppendorf tube. Click chemistry was initiated by adding a 20 µL premixed solution of 1:1 Cu:TBTA (10 mM, prepared from 1 mg CuBr + 0.7 mL 10 mM TBTA in 4:3:1 water:DMSO:t-BuOH). The cap of the tube was removed, and the tube was immediately placed in a 20 mL vial equipped with a rubber septum, followed by Ar flushing for 5 min. We incubated the sealed vial in the dark for two hours. To this DNA solution, we added 10 µL of 3 M sodium acetate (pH 5.2) and 330 µL of 100% ethanol, followed by freezing at -80 °C for 30 min. The frozen stock was then centrifuged for 30 min at 21,000 × *g* at 4 °C to precipitate the DNA. We resuspended the material in 350 µL of 1X bind and wash buffer (B&W; 5 mM Tris, 0.5 mM EDTA, 1 M NaCl, pH 7.5).

We then added 350 µL MyOne C1 streptavidin beads to a 1.5 mL Eppendorf tube. We captured the beads on the side of the tube with a magnet and removed the supernatant. The beads were washed three times with 350 µL 1X B&W. The click product was added to the beads and mixed on a rotator for 30 min. The beads were then captured on a magnetic rack and the supernatant was discarded. The beads were washed three times with 350 µL 1X B&W, and then treated with 100 µL freshly-prepared 0.25 M NaOH solution to generate single-stranded DNA (ssDNA). The beads were captured by magnet, and the supernatant was collected and desalted using a Centri-Sep column (Princeton Separations).

We deprotected the acetyl groups by adding 200 µL concentrated ammonium hydroxide (18 M) to the collected oligos and incubating for 4 hours at room temperature. 450 µL n-butanol was then added to the solution, followed by vortexing, and centrifuging at 21,000 × *g* at 4 °C for 1 min. The top organic layer was removed and discarded. The resulting base-modified aptamer solution was then desalted by a Centri-Spin-10 column (Princeton Separations).

**General procedure for generating particle-displayed base-modified aptamers**

Monoclonal, particle-displayed base-modified aptamers were generated by emulsion PCR. The oil phase was made up of 4.5% Span 80, 0.45% Tween 80, and 0.05% Triton X-100 in mineral oil. The aqueous phase consisted of 1x KOD XL DNA polymerase buffer, 50 U KOD XL DNA polymerase, 0.2 mM dATP, 0.2 mM dGTP, 0.2 mM dCTP (or 0.2 mM 5-formyl-deoxycytidine for the Con A screen), 0.2 mM C8-alkyne-dUTP or DBCO dUTP, 10 nM FP (**C-FP** or **E-FP**), 1 µM fluorescently-labeled RP (**C-RP** or **E-RP**), ~1 pM template DNA, and ~$10^8$ 1 µm FP-conjugated magnetic beads. For each reaction, 1 mL of aqueous phase was added to 7 mL of oil phase and emulsified at 620 rpm for 5 min in an IKA DT-20 tube using the IKA Ultra-Turrax device. The emulsion was pipetted into 100 µL reactions in a 96-well plate. PCR conditions for the Con A library were 96 °C for 2 min; 25 cycles of 96 °C for 15s, 52 °C for 30s, 75 °C for 30s; and 75 °C for 5 min. For the epinephrine library, PCR conditions were 96 °C for 2 min; 25 cycles of 96 °C for 15s, 57 °C for 30s, 74 °C for 30s; and 74 °C for 5 min.

After PCR, the emulsions were collected into an emulsion collection tray (Life Technologies) by centrifuging at 300 x g for 2 min. The emulsion was broken by adding 10 mL 2-butanol to the tray, and the sample was transferred to a 50 mL tube. The tube was vortexed for 30s, and the particles were pelleted by centrifugation at 3,000 x g for 5 min. The oil phase was carefully removed, and the particles were resuspended in 1 mL of emulsion breaking buffer (100 mM NaCl, 1% Triton X-100, 10 mM Tris-HCl, pH 7.5, and 1 mM EDTA) and transferred to a new 1.5 mL tube. After vortexing for 30s and 90s of centrifugation at 15,000 × g, the supernatant was removed. The tube was placed on a magnetic separator (MPC-S, Life Technologies), and the remaining supernatant was removed. The particles were washed three times with 1x PBS buffer using magnetic separation, then stored in 200 µL 1x PBS, 0.1% Tween 20 at 4 °C.

*SPAAC:* This click chemistry reaction was employed when p-AMPBA was used as the click modification for the epinephrine selection. ~$10^7$ particles displaying DNA incorporating DBCO-PEG4-dUTP were incubated overnight at room temperature with an excess of p-AMPBA (>1.5 mg) in 1 ml of storage buffer (10 mM Tris, pH 7.4, 0.025% Tween 20). Before use in the particle display binding assays, particles with the p-AMPBA modification were washed two times with 0.5M NaOH for 10 minutes to deprotect the functional group.

*CuAAC*: This click chemistry reaction was employed when mannose was used as the click modification for the Con A selection. ~$10^7$ particles displaying the DNA library incorporating C8-alkyne-dUTP were resuspended in 10 µL 1x PBS and combined with 25 µL of 20 mM $Na_2HPO_4$, pH 7.3 (degassed 15 min with $N_2$) and 5 µL 10% Tween 20 in a 1.5 mL Eppendorf tube. The click reaction was initiated by the addition of 5 µL 2-azidoethyl 2,3,4,6-tetra-O-acetyl-α-D-mannopyranoside (AeMan, 100 mM in methanol) and 2.5 µL of premixed solution of Cu:TBTA (10 mM, 1 mg Cu(I)Br + 10 mM TBTA in 3:1 DMSO:tBuOH). The reaction was vortexed briefly, placed in a 20 mL vial with a septum, flushed with $N_2$ for 5 min, and incubated in the dark with constant vortexing for 2 hours. The reaction tube was placed on the magnetic separator, and the supernatant was removed. The particles were washed five times with 50 µL TE buffer.

For both click-chemistry procedures, we generated ssDNA by resuspending the particles in 200 µL 0.1 M NaOH solution and incubating for 5 min at room temperature. For the epinephrine screen, particles were washed five times with TE buffer and resuspended in 200 µL 10 mM Tris, pH 7.5. For the Con A screen, the particles were resuspended in 200 µL concentrated ammonium hydroxide (18 M) to deprotect the AeMan. The particles were then incubated for three hours on a slow rotator, washed five times with TE buffer, and resuspended in 200 µL 10 mM Tris, pH 7.5.

We confirmed that base-modified aptamers are efficiently displayed on the particle surface by fluorescently labeling the 3'-end of the base-modified aptamers to allow for FACS characterization after emulsion PCR (**Figure S5**). A cleavable disulfide linker was also incorporated between the aptamer and the particle to allow cleavage of the modified DNA for electrophoretic analysis (**Figure S6**). The slightly lower mobility of the cleaved base-modified aptamer was attributed to the extra mass from the "scar" of the disulfide linker.

**Screening polymerases for efficient reverse transcription**

We optimized the 'reverse transcription' process to convert the carbohydrate-modified DNA back to natural DNA molecules with the same nucleotide sequence. Base-modified aptamer-displaying particles were subjected to PCR under the following conditions: 1 X polymerase buffer,

0.2 mM dATP, 0.2 mM dGTP, 0.2 mM dCTP, 0.2 mM dTTP, 0.4 μM **T-FP**, 0.4 μM **T-RP**, 0.05 U/μL DNA polymerase, $10^4$ base-modified aptamer (**M1**)-displaying particles, and water in a total volume of 50 μL. PCR conditions were as follows: 96 °C for 2 min; 30 cycles of 96 °C for 15 s, 51 °C for 30 s, and 72 °C for 30 s; 72 °C for 2 min; and hold at 4 °C. We screened four DNA polymerases—Taq, KOD-XL, Pwo, and Deep Vent—for the efficiency of reverse transcription. 2 μL of each PCR reaction was loaded directly onto a 10% native PAGE gel and run at 150 V for 30 min in 1X TBE buffer. Gels were imaged after staining with 1X GelStar Nucleic Acid Stain in TBE buffer. After testing several DNA polymerases (**Figure S7a**), we found that Taq efficiently generated DNA of the correct length from the base-modified aptamers (**Figure S7b**). Sanger sequencing showed that the product generated by Taq polymerase was identical to the starting template, confirming the fidelity of the reverse transcription process (**Figure S8**).

The same experimental procedure was performed for base-modified aptamer **M1**-displaying particles with the boronic acid modification, and we determined that Taq polymerase efficiently reverse-transcribed our sequence. (data not shown)

**Pre-enrichment of the DNA library for epinephrine**

Norepinephrine and epinephrine differ by a single methyl group. Since we are targeting the diol structure on the molecule, we used the amine handle on norepinephrine to conjugate the molecule to magnetics beads.

500 ul MyOne COOH beads were washed three times with DMSO. 13.8 mg N-hydroxysuccinimide in 250 μl DMSO, 19.1 mg 1-ethyl-3-(3-dimethylaminopropyl)carbodiimide hydrochloride in 250 μl DMSO and 30 μl triethylamine were added to the beads and incubated with rotation for 30 minutes. Beads were washed with 1 mL DMSO. 1 mL 100 mg/mL norepinephrine in DMSO was incubated with beads for 3 hr at room temperature with rotation. After conjugation, beads were washed five times with DMSO, and five times with 1X PBS, 100 mM NaCl.

6 nmol random library (Primers: **E FP**, **E RP**, **Table S1,** with an N40 random region) was incubated with 100 μl epinephrine conjugated beads in a total of 200 μl selection buffer (20 mM Tris, pH 7.5, 120 mM NaCl, 5 mM KCl, 1 mM $MgCl_2$, 1 mM $CaCl_2$, 0.1% Tween 20) for 30 minutes at room temperature, with rotation. Beads were washed once with 200 μl selection buffer. DNA was eluted from the beads with 200 μl 0.5 M NaOH. Beads were washed once more with 50 μl 0.1 M NaOH. DNA was recovered from NaOH by adjusting the pH with 25 μl 3M NaOAc, then purified with a Qiagen MiniElute cleanup kit. DNA was amplified with biotinylated reverse primer **E-RP**. PCR conditions were 96 °C for 2 min; 25 cycles of 96 °C for 15s, 57 °C for 30s, 74 °C for 30s; and 74 °C for 5 min.

ssDNA was generated by using MyOne SA C1 beads to capture the double-stranded DNA, after which the sense strand was eluted by incubating the DNA-coated beads with 0.1 M NaOH for 10 minutes. DNA was recovered by adding 1/10 vol 3M NaOAc to neutralize pH followed by purification with a Qiagen MiniElute cleanup kit.

SELEX was repeated with the isolated sense strands under the same conditions (100ul epinephrine beads in 200ul selection buffer, rotating 30 min RT) and recovered in the same way. The output DNA of the second round of SELEX was used as the input DNA for particle display

**Preparation of FITC-epinephrine conjugate**

7.35 μmol FITC-PEG (3,000 Da)-NHS and 73.5 μmol epinephrine were combined in 1 mL 1x PBS (which had previously been bubbled in $N_2$ for 10 min) and 2 mL DMSO, and then incubated overnight with rotation at room temperature. The FITC-epinephrine conjugate was purified by adding 40 mL 1x PBS and spinning through 3k spin filters (Amicon) until the flow-through was clear. The conjugate was concentrated to 1 mL in the spin filter. Unreacted NHS was removed by incubating with 200 μl amine 270-μm Dynabeads (Thermo Fisher) for 3 hours at room temperature. Supernatant was saved for particle display selection by magnetic separation.

**Click-PD screening**

For each round of screening for epinephrine, ~$10^7$ boronic acid-modified aptamer particles were incubated in 100 ul binding buffer at 95 °C for 2 min, and then cooled at room temperature for 30 min. The beads were incubated with the appropriate concentration of FITC-epinephrine conjugates in 1mL on the rotator in the dark for 1 h. After removing the supernatant, the particles were resuspended in 1 mL selection buffer, and then analyzed using a BD FACS Aria III. In each round, the sort gate was set to collect aptamer particles that show high binding affinity towards epinephrine (*i.e.*, high FITC fluorescence). As the rounds progressed, we applied higher stringency for the sort gates. The collected population ranged from 0.1–0.3%. After sorting, the collected base-modified aptamer particles were resuspended in 20 μL PBS and reverse transcribed into canonical DNA by Taq polymerase.

For each round of screening for Con A, we incubated ~$10^8$ mannose-modified aptamer particles with 1 nM biotinylated Con A (Sigma) and 250 nM FITC-conjugated PSA (Sigma) in selection buffer (SB; 1 x PBS, 2.5 mM $MgCl_2$, 1 mM $CaCl_2$, 0.1 mM $MnCl_2$, 0.01% Tween 20) for 1 hour in the dark on a rotator. After incubation, 2 μL of 2 mg/mL streptavidin-conjugated Alexa Fluor 647 was diluted in 998 μL of selection buffer. The aptamer particles were resuspended in the 1 mL mixture and incubated for 10 min in the dark on a rotator to fluorescently label the biotinylated Con A. The particles were washed once and resuspended in SB. The sample was then analyzed with the BD FACS Aria III, with the sort gate set to collect base-modified aptamer particles that exhibit high binding to Con A and low binding to PSA. 0.2–1.0% of the total singlet population was collected in each round. After sorting, the collected base-modified aptamer particles were resuspended in 20 μL PBS and reverse transcribed into canonical DNA by Taq polymerase.

For both targets, collected aptamer particles were subjected to PCR for the next round with 1X Taq PCR Mastermix, 40 nM FP, 40 nM RP, and nuclease-free water in a volume of 50 μL. PCR conditions for the Con A library were 96 °C for 2 min; 25 cycles of 96 °C for 15s, 52 °C for 30s, 75 °C for 30s; and 75 °C for 5 min. For the epinephrine library, PCR conditions were 96 °C for 2 min; 25 cycles of 96 °C for 15s, 57 °C for 30s, 74 °C for 30s; and 74 °C for 5 min.

After 10 rounds of amplification, further cycle optimization was carried out to generate the DNA library for the next round. Products from additional cycles of PCR were analyzed via a 10% native PAGE gel, run at 170 V for 40 min in 1X TBE buffer. After staining the gel with 1X GelStar Nucleic Acid Gel Stain, the gel was imaged by Gel-Doc (Bio-Rad). The optimized cycle number was used to complete DNA amplification with the remaining aptamer particles (1X Taq PCR Mastermix, 40 nM FP, 40 nM RP, and nuclease-free water in a final volume of 400 μL). The final PCR reaction was cleaned up using a Qiagen MinElute Reaction Cleanup kit.

**High-throughput sequencing of the enriched libraries**

DNA pools for high-throughput sequencing were prepared as described in *16S Metagenomic Sequencing Library Preparation* by Illumina (https://support.illumina.com/downloads/16s_metagenomic_sequencing_library_preparation.html). Overhang adaptor sequences for the forward and reverse primers were ordered from IDT. DNA pools from each round were indexed using the Nextera XT DNA Library Preparation Kit (Illumina) and then pooled for sequencing. Sequencing was performed using an Illumina MiSeq at the Stanford Functional Genomics Facility. FASTQ data were analyzed using Galaxy NGS tools. Read 1 and Read 2 datasets were merged via the 'Pear' program, and then trimmed via 'Trim sequences'. Sequences with low quality were filtered out using "Filter by quality", accepting only sequences with more than 90% of the bases having a quality score of 20 or above. The FASTAptamer toolkit was used to identify sequence clusters (sequences varying by 2 or fewer bases) and calculate the degree enrichment of each sequence from round to round.

**General procedure for particle-based binding assay for fluorescently-labeled targets**

~$10^6$ particles were incubated with varying concentrations of fluorescently-labeled protein in SB for 1 hour on a rotator at RT. After incubation, the particles were washed once in 100 uL cold SB and resuspended in 100uL cold SB. The particles were analyzed using the BD Accuri C6 flow cytometer, and the mean fluorescence and/or percentage of bound particles were measured in the relevant fluorescence channel(s).

**Generation of solution-phase base-modified aptamers with 5'-biotinylation for ConA3-1m**

Base-modified aptamers were amplified in a PCR mixture containing 1X KOD-XL polymerase buffer, 0.2 mM dATP, 0.2 mM dGTP, 0.2 mM 5-formyl-deoxycytidine, 0.2 mM C8-alkyne-dUTP, 0.4 μM 5'-biotinylated **C-FP-Bio**, 0.4 μM **C-RP**, 0.05 U/μL KOD-XL DNA polymerase, 20 pM PCR template, and water in a total volume of 5 mL in a 96 well plate. PCR conditions were as follows: 96 °C for 2 min; 12 cycles of 96 °C for 15 s, 52 °C for 30 s, and 75 °C for 30 s; 75 °C for 2 min, and hold at 4 °C.

PCR reactions were transferred into a 50 mL conical tube. To this PCR mixture, we added 0.5 mL 3 M sodium acetate (pH 5.2) and 13.75 mL of 100% ethanol, followed by freezing at -80 °C for 30 min. The frozen stock was then centrifuged for 30 min at 4000 x g at 4 °C to precipitate the DNA. The pellet was dissolved with 600 μL water, followed by purification using MinElute spin columns. The PCR product was eluted with 180 μL of 10 mM Tris buffer, pH 8.0. We then added 40 μL of 3 M sodium acetate (pH 5.2) and 1.2 mL of 100% ethanol, followed by freezing at -80 °C for 30 min. The frozen stock was then centrifuged for 30 min at 21,000 × *g* at 4 °C to precipitate the DNA. The DNA was resuspended in 20 μL 1X PBS buffer.

We combined 20 μL of the base-modified DNA solution with 20 μL of 100 mM 2-azidoethyl 2,3,4,6-tetra-O-acetyl-α-D-mannopyranoside in DMSO (100 eq) and 40 μL 20 mM sodium phosphate buffer, pH 8 (pre-degassed by bubbling $N_2$ through) in a 1.5 mL Eppendorf tube. Click chemistry was initiated by the addition of 20 μL of a premixed solution of 1:1 Cu:TBTA (10 mM, prepared with 1 mg CuBr + 0.7 mL of 10 mM TBTA in 4:3:1 water:DMSO:t-BuOH). The cap of the tube was removed, and the de-capped tube was immediately placed in a 20 mL vial equipped with a rubber septum, followed by Ar flushing for 5 min. We incubated the sealed vial in the dark for two hours. To this DNA solution, we added 10 μL of 3 M sodium acetate (pH 5.2) and 330 μL of 100% ethanol, followed by freezing at -80 °C for 30 min. The frozen stock was then centrifuged for 30 min at 21,000 ×*g* at 4 °C to precipitate the DNA. We resuspended the DNA in 350 μL 1X B&W.

We added 350 μL MyOne C1 streptavidin beads to a 1.5 mL Eppendorf tube. The beads were captured on the side of the tube with a magnet and the supernatant was removed. The beads were washed three times with 350 μL 1X B&W. The click product sample was added to the beads and mixed on a rotator at room temperature for 30 min. The beads were then captured and the supernatant was discarded. The beads were washed three times with 350 μL 1X B&W, then treated twice with 100 μL of freshly-prepared 0.25 M NaOH solution to generate ssDNA. The supernatant was discarded. The acetyl group on the mannose was deprotected by adding 300 μL of concentrated ammonium hydroxide (18 M) and incubating at room temperature for three hours. This tube was then sealed tightly before heating on a thermal block at 70 °C for 10 min. The sample was cooled in an ice bath before opening the cap. The tube was placed on the magnet, and the supernatant was transferred to a separate tube. 100 μL of 18 M ammonium hydroxide was again added to the beads, and the heating procedure was repeated.

The supernatants from the two ammonium hydroxide treatment steps were combined and then mixed with 4.5 mL n-butanol before vortexing and centrifuging at 16,000 ×*g* at 4 °C for 10 min. The supernatant was removed and discarded. The sample was dried over vacuum centrifugation, and then resuspended in 100 μL water. To this solution, we added 50 μL of 5 M $NH_4OAc$ and 415 μL of cold 100% ethanol before freezing at -80 °C for 30 min. We centrifuged for 30 min at 21,000 ×*g* at 4 °C to precipitate the base-modified aptamer. The pellet was washed once with 70% cold ethanol in water, then dissolved in 100 μL water.

**Bio-layer interferometry measurement of selected base-modified aptamers**

**ConA-3-1** and **ConA-3-1m** were diluted to 50 nM in SB. Solutions of 0, 1, 2, 4, 8, 16, 32, and 64 nM Con A were prepared in SB. The solutions were loaded into a 384-well plate, with 100 μL of SB, 80 μL of biotinylated aptamer, and 100 μL of Con A solution for each reaction. The following steps were run on the ForteBIO Octet RED384 with Super Streptavidin biosensors: 60s in buffer for equilibration, 5 min in aptamer solution to load the aptamer onto the biosensors, 60s in buffer for a baseline measurement, 10 min in Con A solution to measure association, and 10 minutes in buffer to measure dissociation. Analysis was performed using Octet Data Analysis software, including the alignment of the different measurements and global fitting of the experimental data to a binding model to extract $K_d$, $k_{on}$, and $k_{off}$.

**Lectin array assay to probe base-modified aptamer specificity**

The following procedure was adapted from RayBiotech's product manual for the Lectin Array 70. First, we dried the glass slide. The slide with the pre-printed lectin array was equilibrated to room temperature inside the sealed plastic bag for 20–30 minutes. We then annealed 30 μL of 0.5 μM **ConA-3-1m** in 1X PBS by incubating the solution at 95 °C and slowly cooling down to 4 °C at a ramp rate of 0.1 °C/second. We incubated at 4 °C for 5 min. We then added 100 μL sample diluent (included in the lectin array package) into each well of the array and incubated at room temperature for 30 min to block the slides. We removed the buffer from each well. After diluting **ConA-3-1m** to the desired concentration with SB, we added 100 μL of diluted **ConA-3-1m** to each well and incubated the arrays at room temperature for 3 hours. We then removed the samples from each well, and washed each well five times (5 min each) with 150 μL of 1X wash buffer I (included in the lectin array package, supplemented with 2.5 mM $MgCl_2$, 1 mM $CaCl_2$, and 0.1

mM $MnCl_2$) at room temperature with gentle shaking. We completely removed the buffer between each wash step. We then washed two times (5 min each) with 150 μL of 1X wash buffer II (included in the lectin array package, supplemented with 2.5 mM $MgCl_2$, 1 mM $CaCl_2$, and 0.1 mM $MnCl_2$) at room temperature with gentle shaking. We completely removed the wash buffer between each wash step. We then briefly spun down the Cy3 equivalent dye-conjugated streptavidin tube (included in the lectin array package), and added 1.4 mL of sample diluent to the tube, mixing gently. We added 80 μL of Cy3 equivalent dye-conjugated streptavidin to each well and incubated in the dark at room temperature for 1 hour. We decanted the samples from each well, and washed five times with 150 μL of 1X wash buffer I at room temperature with gentle shaking, completely removing the wash buffer after each wash step. We disassembled the slide assembly by pushing the clips outward from the slide side and carefully removing the slide from the gasket. We placed the slide in the slide washer/dryer (a four-slide holder/centrifuge tube included in the lectin array package), adding enough 1x wash buffer I to cover the whole slide (~30 mL), and then gently agitated at room temperature for 15 minutes. After decanting wash buffer I, we washed with 1x wash buffer II (about 30 mL) with gentle shaking at room temperature for 5 minutes. Finally, we dried the slide by centrifugation at 200 × *g* on a microscope slide spinner and scanned the slide on a GenePix μArray scanner, monitoring the Cy3 dye channel at PMT 500.

**Determining Con A concentration to induce complete hemagglutination**

Human erythrocytes were washed and resuspended in 1X PBS in a 96-well U-shaped well plate at 1% hematocrit, with Con A concentrations ranging from 2–250 μg/mL. We let the plate stand at room temperature for 1 hour before visualizing the deposition of erythrocytes at the bottom of the well. The optical densities at 655 nm were then measured on a Tecan M220 plate reader.

**Hemagglutination inhibition assay**

We annealed 30 μL of 0.5 μM **ConA-3-1m** in 1X PBS by heating the solution to 95 °C and slowly cooling down to 4 °C at a ramp rate of 0.1 °C/second, followed by incubation at 4 °C for 5 min. We incubated the annealed aptamer at a range of concentrations from 9.6–300 nM with 150 nM Con A in 1X PBS for 30 min in a 96-well U-shaped well plate. Human erythrocytes were added to produce a cell suspension of 1% hematocrit in a total volume of 50 μL per well. After 1 hour of incubation at RT, the hemagglutination status of the samples was visualized, and the optical densities of the cell suspensions at 655 nm were monitored by a plate reader.

**Microscopic characterization of human erythrocyte agglutination**

We annealed 2 μL of 4 μM **ConA-3-1m** in 1X PBS by incubating the solution at 95 °C and slowly cooling down to 4 °C at a ramp rate of 0.1 °C/second, and then incubated at 4 °C for 5 min. We then added 1 μL of 6.5 μM Con A and incubated for 30 min at room temperature. We prepared an erythrocyte suspension to a final hematocrit of 20% in PBS, and 7 μL of erythrocyte suspension was combined either with the Con A-aptamer complex or 3 μL of 1X PBS. 10 μL of this mixture was loaded onto glass slides, covered with coverslips, and immediately visualized using 10X and 40X objective lenses on a microscope. Optical microscopy imaging was performed on an Olympus CKX-41 inverted microscope with color digital camera. The images were processed with ImageJ software.

**Table S1. DNA sequences used in this study.**

| **Name** | **DNA sequence** |
|---|---|
| **S1** | 5'-CGG AAC GTC /i5OctdU//i5OctdU//i5OctdU/ GTA ACT TGA-3' |
| **T1** | 5'- ATC CAG AGT GAC GCA GCA CGG AAC GTC TTT GTA ACT TGA AAT ACC GTG GTA GGT TGG CTA GGT TGG ACA CGG TGG CTT AGT -3' |
| **M1** | 5'- ATC CAG AGT GAC GCA GCA **2**GG AA**2** G**42** **444** G**4**A A**24** **4**GA AA**4** A**22** G**4**G G**4**A GG**4** **4**GG **24**A GG**4** **4**GG A**2**A **2**GG **4**GG **244** AG**4** -3' |
| **T-FP** | 5'- ATC CAG AGT GAC GCA GCA -3' |
| **T-RP** | 5'- ACT AAG CCA CCG TGT CCA -3' |
| **T-RP-2Bio** | 5'- /52-Bio/ACT AAG CCA CCG TGT CCA -3' |
| **T-29-DBCO** | CGGAACGTC**4**GTAACTTGA |
| **Epinephrine library and test sequences** | |
| **Library** | 5'- **E-FP--**N(40)--**E-RPcomp**-3' |
| **E-FP** | 5'- CCA GCG AGC CAG CGA C -3' |
| **E-RP** | 5'- CTG TGC CGT CCT GCG TG -3' |
| **4-1** | 5'- CCA GCG AGC CAG CGA CG/DBCO/ ACG /DBCO/GA A/DBCO/C CA/DBCO/ GGG GAC GGA GAG ACG GCA CGA ACC AAC ACG CAG GAC GGC ACA G -3' |
| **ConA library and test sequences** | |
| **Library** | 5'- **C-FP--**N(40)--**C-RPcomp**-3' |
| **C FP** | 5'- GAT CCC AGT CCG AAG TAA TC -3' |
| **C RP** | 5'- CCT ATA GCC GTT TGC ACA AG -3' |
| **ConA Seq 1** | 5'- GAC AAG GAA AAT CCT TTC AAT GAA GTG GGT C -3' |
| **ConA-3-1** | 5'-GAT CCC AGT CCG AAG TAA TCG **44**G **2**A**4** **24**G **2**A**2** GA**2** **4**GG **4**GA G**24** **4**GA G**4**G G**2**A GAA GAA **244** G**4**G **2**AA A**2**G G**24** A**4**A GG-3' |
| **ConA-3-1m** | 5'-GAT CCC AGT CCG AAG TAA TCG **44**G CA**4** C**4**G CAC GAC **4**GG **4**GA GC**4** **4**GA G**4**G GCA GAA GAA C**44** G**4**G CAA ACG GC**4** A**4**A GG-3' |
| **ConA-3-1a** | 5'-GAT CCC AGT CCG AAG TAA TCG TTG **2**AT **2**TG **2**A**2** GA**2** TGG TGA G**2**T TGA GTG G**2**A GAA GAA **2**TT GTG **2**AA A**2**G G**2**T ATA GG-3' |

| | |
|---|---|
| **ConA-3-1nc** | 5'-GAT CCC AGT CCG AAG TAA TCG **11**G **2**A**1** **21**G **2**A**2** GA**2** **1**GG **1**GA G**21** **1**GA G**1**G G**2**A GAA GAA **211** G**1**G **2**AA A**2**G G**21** A**1**A GG-3' |
| **ConA-3-1n** | 5'-GAT CCC AGT CCG AAG TAA TCG TTG CAT CTG CAC GAC TGG TGA GCT TGA GTG GCA GAA GAA CTT GTG CAA ACG GCT ATA GG-3' |
| **ConA-3-1mscr** | 5'- GAT CCC AGT CCG AAG TAA TCA GCA G**44** AA**4** G**44** AGG A**4**G CGG AGG CGC A**4**A CG**4** CG**4** ACG C**44** G**4**G CAA ACG GC**4** A**4**A GG-3' |

The following modifications were requested when ordering from IDT, using IDT's nomenclature: /i5OctdU/: C8-alkyne-deoxyuridine; /52-Bio/: 5'-dual biotin modifier; /DBCO/: DBCO-dT-CE phosphoramidite (Glen Research, Catalog# 10-1539)

**1**: C8-alkyne-dU; **2**: 5-formyl-dC; **4**: 2-azidoethyl 2,3,4,6-tetra-O-acetyl-α-D-mannopyranoside-modified **1**; See **Figure 3** (main text) for the structures of **1**, **2**, and **4.**

#### II. Supplementary Results

**Figure S1.**

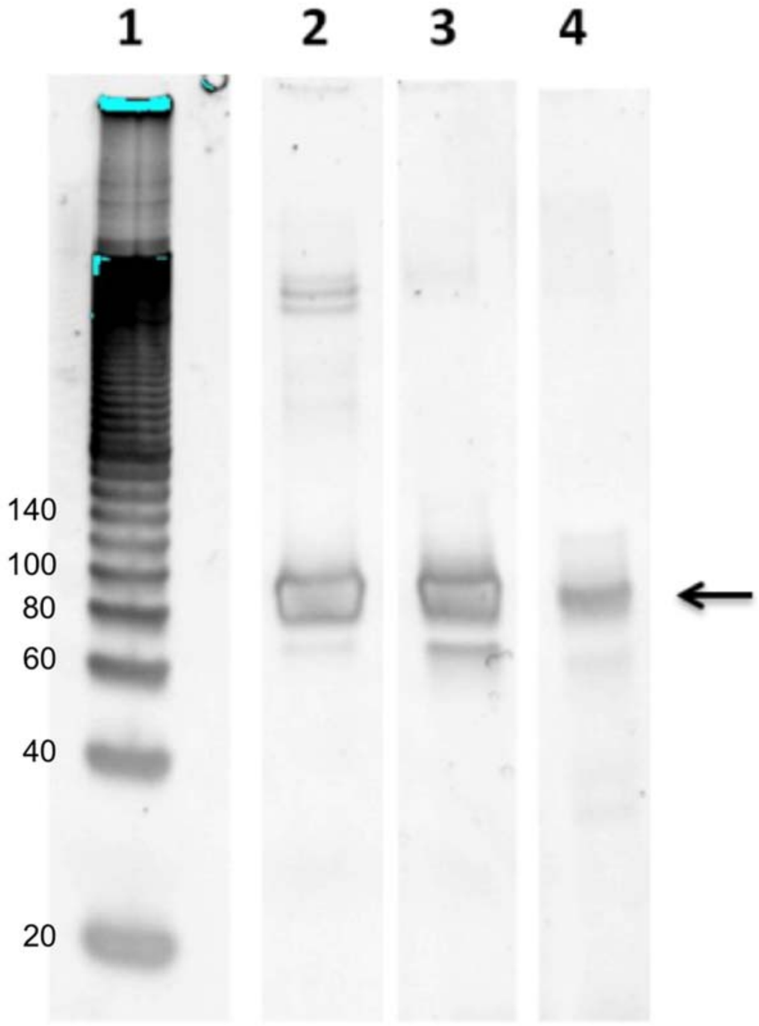

**Figure S1. Screen for polymerase-mediated incorporation of modified pyrimidine deoxyribonucleotides C8-alkyne-dU and 5-formyl-dC.** PCR template is the 81-nt DNA oligonucleotide, **T1.** Lane 1: DNA ladder; lane 2: KOD-XL; lane 3: Pwo; lane 4: Deep Vent. The arrow indicates the full-length product. KOD-XL DNA polymerase gives the highest yield without a major byproduct.

**Figure S2.**

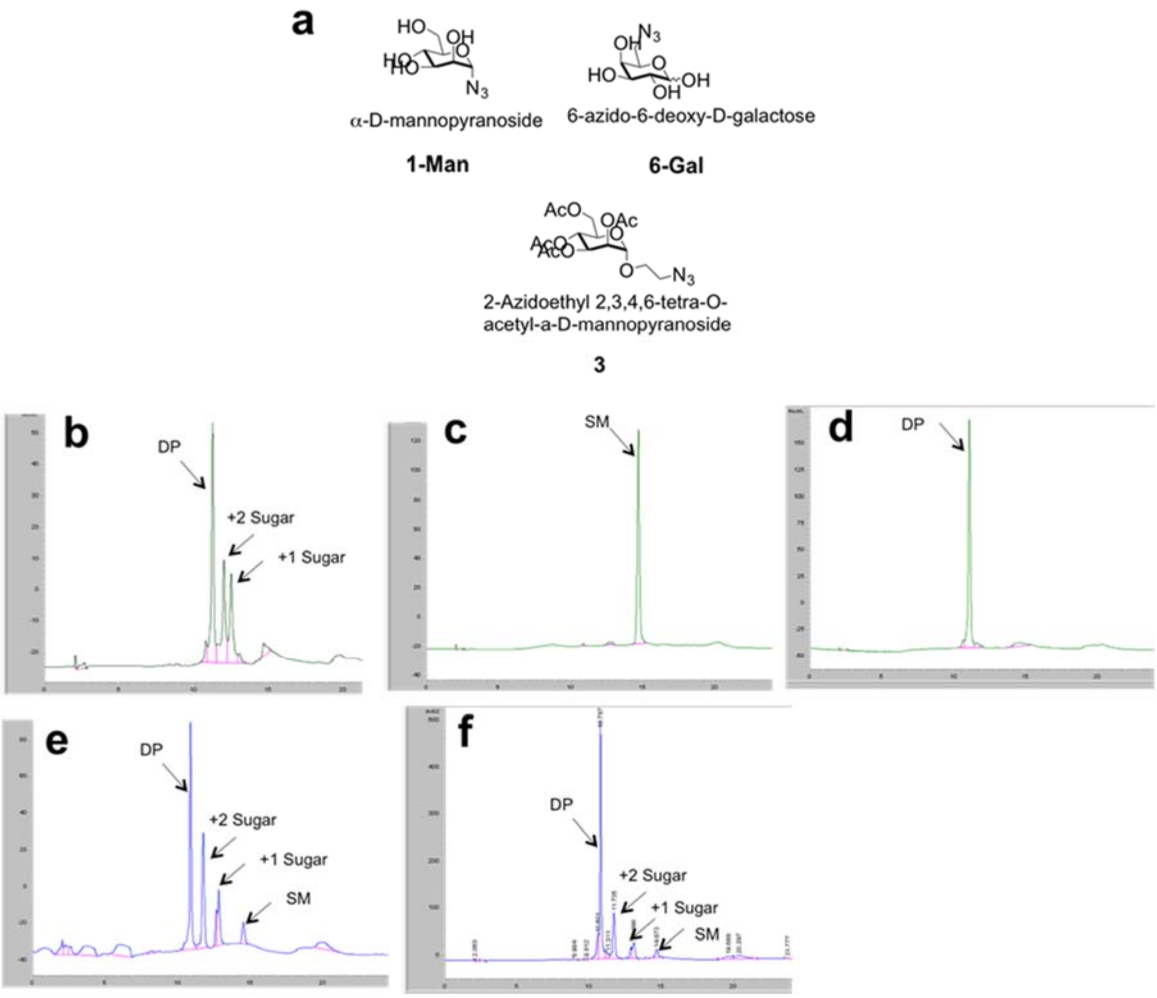

**Figure S2. Optimization of click chemistry using a 21-nt oligonucleotide substrate with three consecutive alkyne side chains.** (a) Structures of azido-sugars screened for click conjugation to an alkyne-bearing 21-nt DNA oligonucleotide with three consecutive alkyne-modified U nucleotides, **S1**. (b)-(f) HPLC analysis of click conjugation under different conditions with different substrates. Click chemistry conditions are as follows: (b) conjugation of **3** with 0.4 mM $CuSO_4$ + 2 mM THPTA + 4 mM sodium ascorbate; (c) conjugation of **3** with 0.4 mM $CuSO_4$ + 0.4 mM TBTA + 0.8 mM TCEP; (d) conjugation of **3** with 0.4 mM CuBr + 0.4 mM TBTA; (e) conjugation of **1-Man** with 0.4 mM CuBr + 0.4 mM TBTA; (f) conjugation of **6-Gal** with 0.4 mM CuBr + 0.4 mM TBTA. DP: desired product. SM: starting material. +1 sugar and +2 sugar: products with one or two carbohydrate substrates conjugated. Only click conjugation of substrates **3** and AeGla (results not shown) with 0.4 mM CuBr + 0.4 mM TBTA gave quantitative yield of the desired product without major byproducts .

**Figure S3.**

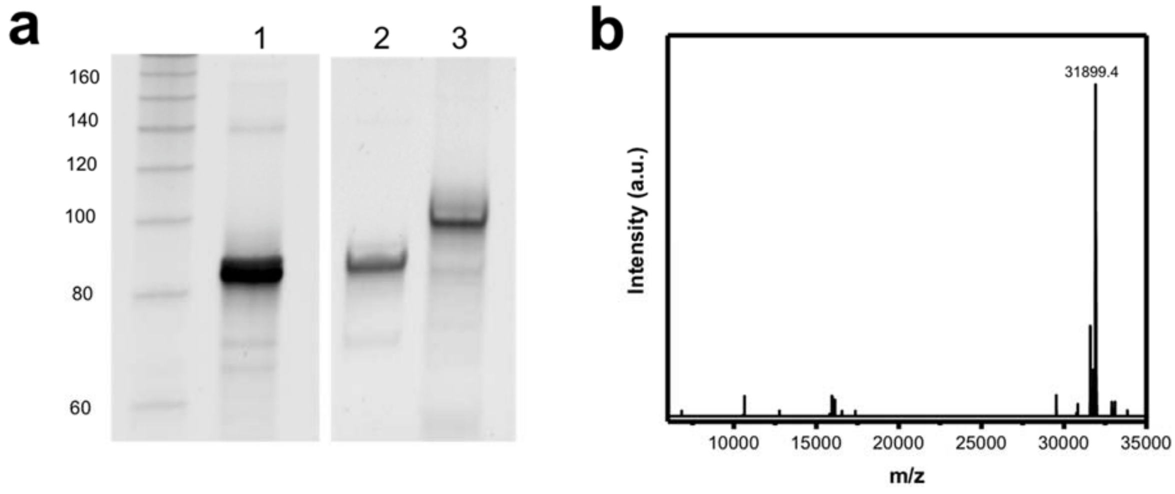

**Figure S3. Optimization of click conjugation.** (a) Our click chemistry reaction efficiently modified the **T1**-derived PCR product (after removing the antisense strand), **M1,** which contains numerous C8-alkyne-dU and 5-formyl-dC nucleotides. Gel lanes represent the template before (lane 1) and after strand separation either immediately after PCR (lane 2) or after subsequent click conjugation with mannose (lane 3). (b) ESI-MS characterization of **M1** (expected: 31901.2 Da, observed: 31899.4 Da).

**Figure S4.**

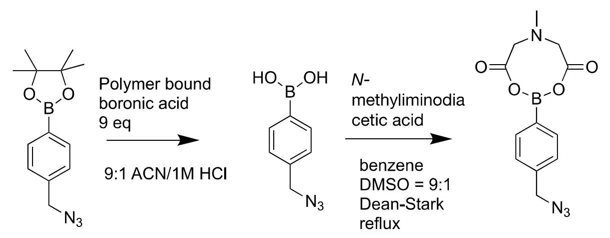

**Figure S4:** synthesis of *N*-methyliminodiacetic acid (MIDA)-protected p-azidomethyl phenylboronic acid (p-AMPBA)

**Figure S5.**

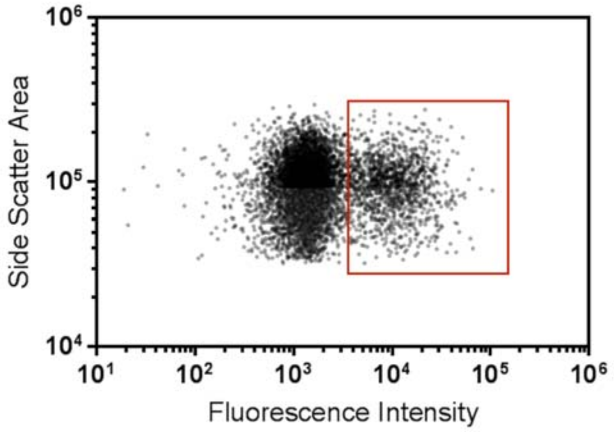

**Figure S5. Characterization of base-modified aptamer particle synthesis using flow cytometry.** Fluorescent signal from the Alexa Fluor 647-labeled reverse primer shows two populations: blank particles, and particles coated with base-modified aptamers (red box). In this representative sample, 18% of the particles are coated with base-modified aptamers.

**Figure S6.**

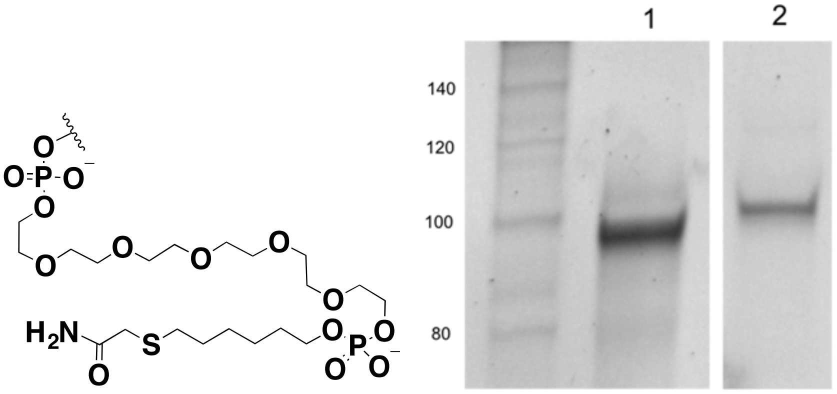

**Figure S6**. **Confirmation of the generation of particle-displayed base-modified aptamers.** Left, structure of the "scar" of the disulfide linker after base-modified aptamer cleavage. The disulfide linker between the forward primer and the particle is cleaved by TCEP treatment followed by alkylation using iodoacetamide. Right, our click chemistry reaction conditions efficiently modified particle-coupled base-modified aptamers. Lane 1 contains the reaction product **M1** formed in solution (see **Figure S3a**), and lane 2 contains base-modified aptamer cleaved from beads after emulsion PCR and on-bead click reaction.

**Figure S7.**

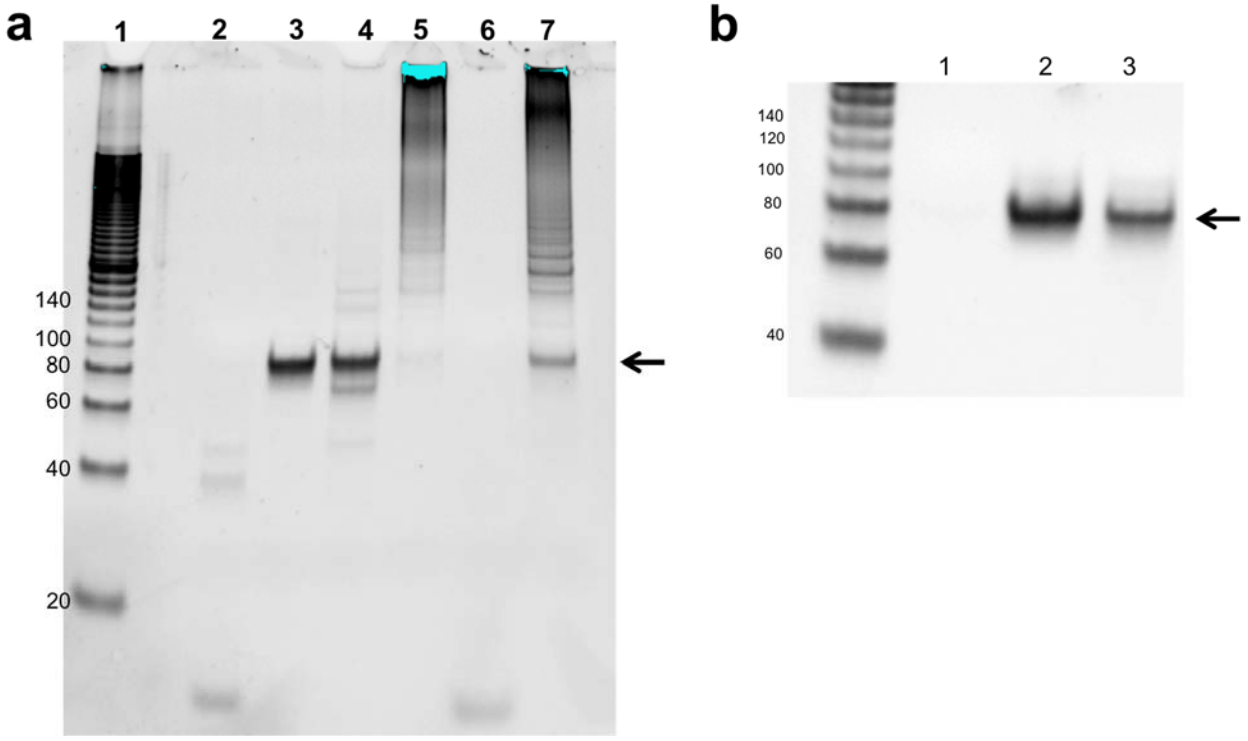

**Figure S7. Taq polymerase efficiently converts base-modified aptamers back to natural DNA via a 'reverse transcription' PCR process.** (a) Polymerase screen for the reverse transcription step. Lane 1: DNA ladder; lane 2: Taq polymerase, without template; lane 3: Taq polymerase, using canonical DNA template **T1**; lane 4: Taq polymerase, using base-modified aptamer particles as template; lane 5: KOD-XL, using base-modified aptamer particles as template; lane 6: Pwo, using base-modified aptamer particles as template; lane 7: Deep Vent, using base-modified aptamer particles as template. The arrow indicates the full-length product. (b) Confirmation of reverse-transcription using Taq DNA polymerase. Lane 1: PCR without template; lane 2: PCR using **T1** as the template; lane 3: PCR using base-modified aptamer **M1** displayed on beads as template.

Polymerases screened were chosen from the following references:

Jager S, Rasched G, Kornreich-Leshem H, Engeser M, Thum O, Famulok M. A versatile toolbox for variable DNA functionalization at high density. *J Am Chem Soc*. 2005;127(43):15071-15082. doi:10.1021/ja051725b.

Hili R, Niu J, Liu DR. DNA ligase-mediated translation of DNA into densely functionalized nucleic acid polymers. *J Am Chem Soc*. 2013;135(1):98-101. doi:10.1021/ja311331m.

**Figure S8.**

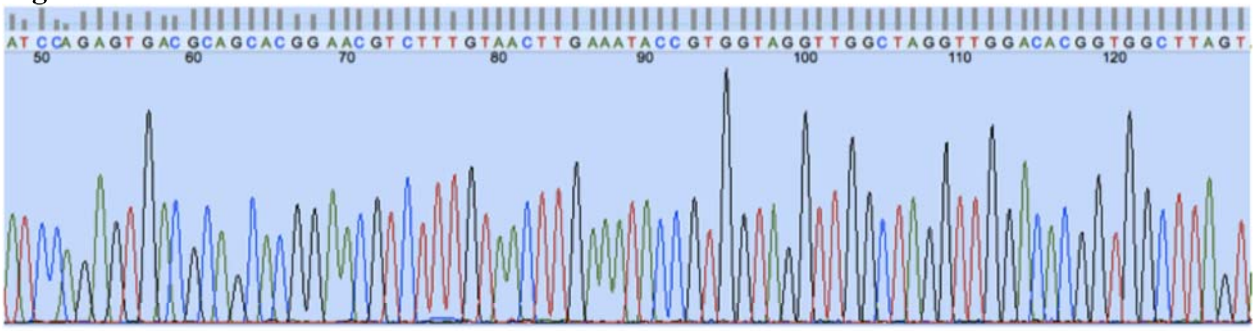

Template used for generating non-natural aptamer-displayed particle: **T1**
5'- ATC CAG AGT GAC GCA GCA CGG AAC GTC TTT GTA ACT TGA AAT ACC GTG GTA GGT TGG CTA GGT TGG ACA CGG TGG CTT AGT -3'

Sequenced PCR product (sense strand) of the reverse transcription:
5'- ATC CAG AGT GAC GCA GCA CGG AAC GTC TTT GTA ACT TGA AAT ACC GTG GTA GGT TGG CTA GGT TGG ACA CGG TGG CTT AGT -3'

**Figure S8. Sanger sequencing of the product of reverse-transcription.** PCR products from reverse transcription were cloned into a TOPO vector and transfected into TOP10 chemically-competent *E. coli*. Colonies were harvested and sent for Sanger sequencing. All 20 colonies sequenced were either matched or complementary to the sequence of **T1**, demonstrating good fidelity for the reverse-transcription reaction.

**Figure S9.**

(a)

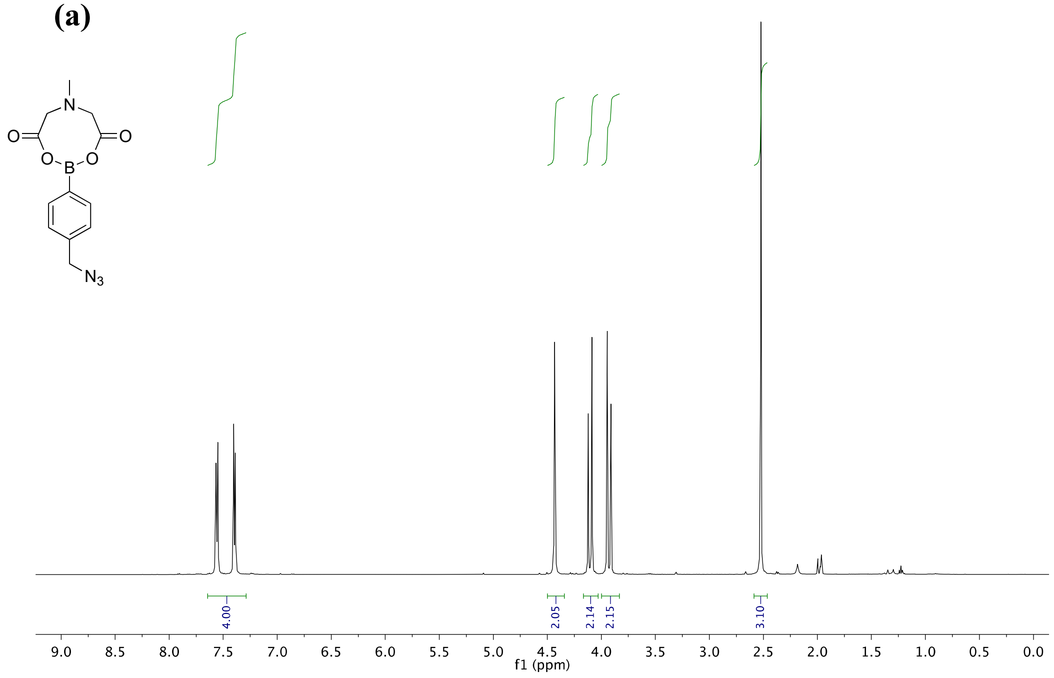

(b)

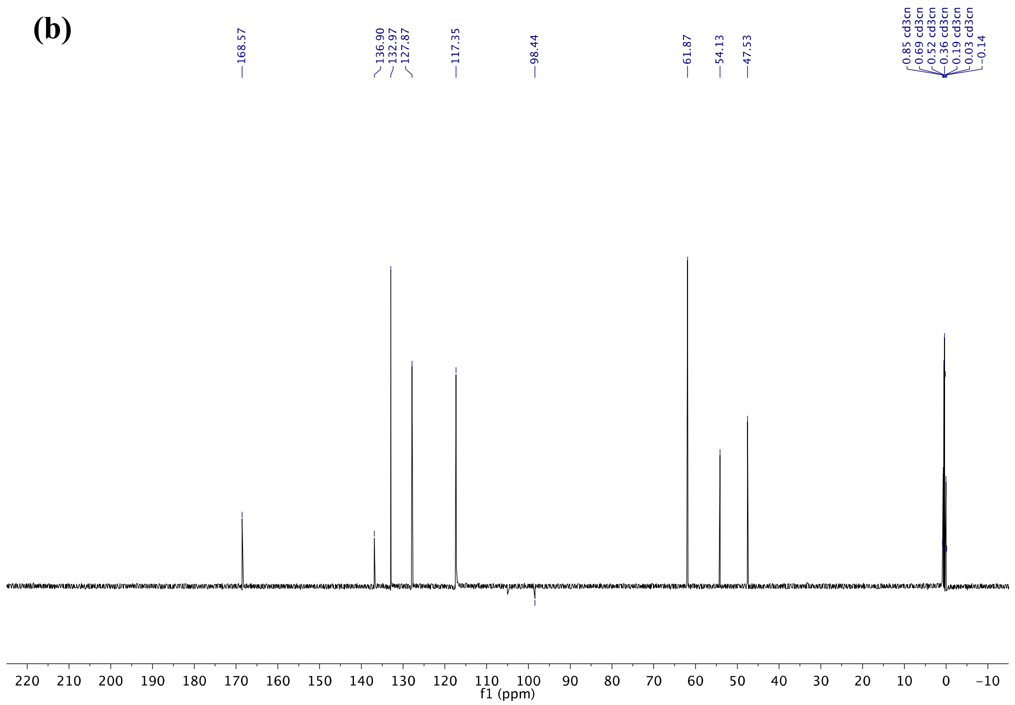

Figure **S9**: $^{1}$H-NMR (**a**) and $^{13}$C-NMR (**b**) spectra of *p*-AMPBA MIDA ester.

**Figure S10.**

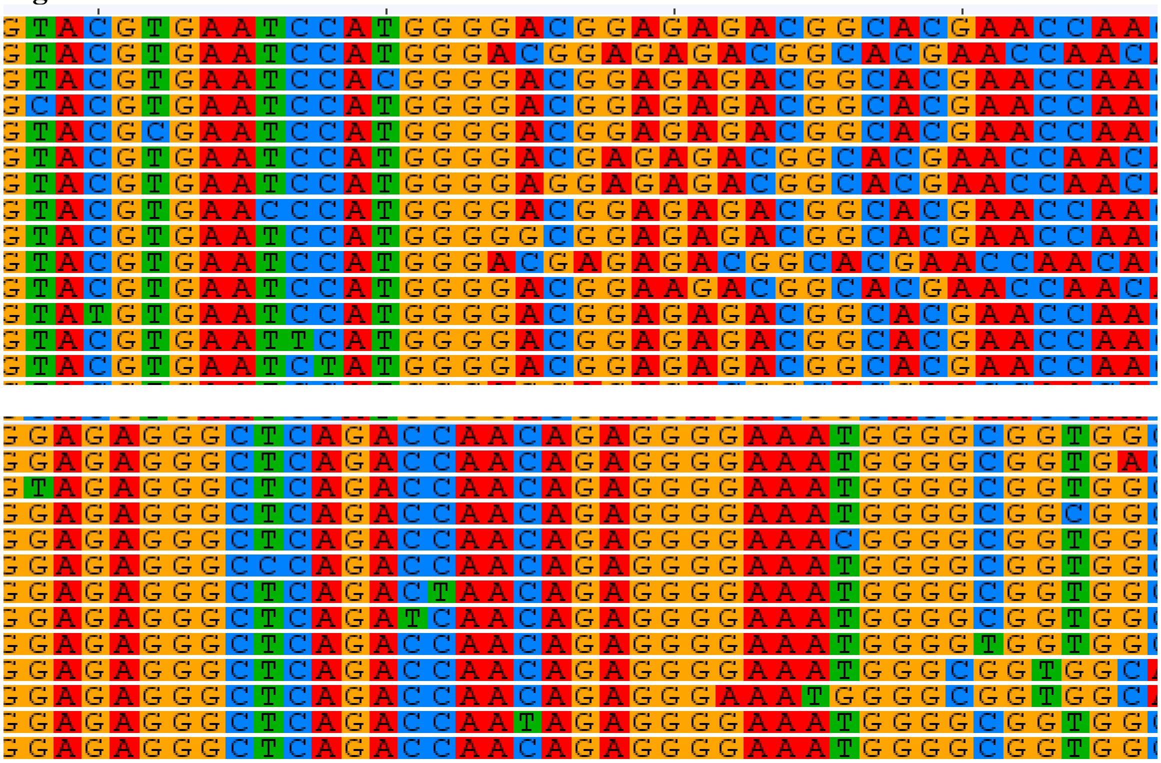

**Figure S10:** Two major families identified from four rounds of click PD for epinephrine. Random region only shown. These families only differ by a shift or single mutation. Two representative sequences from each family were chosen to characterize further.

**Figure S11.**

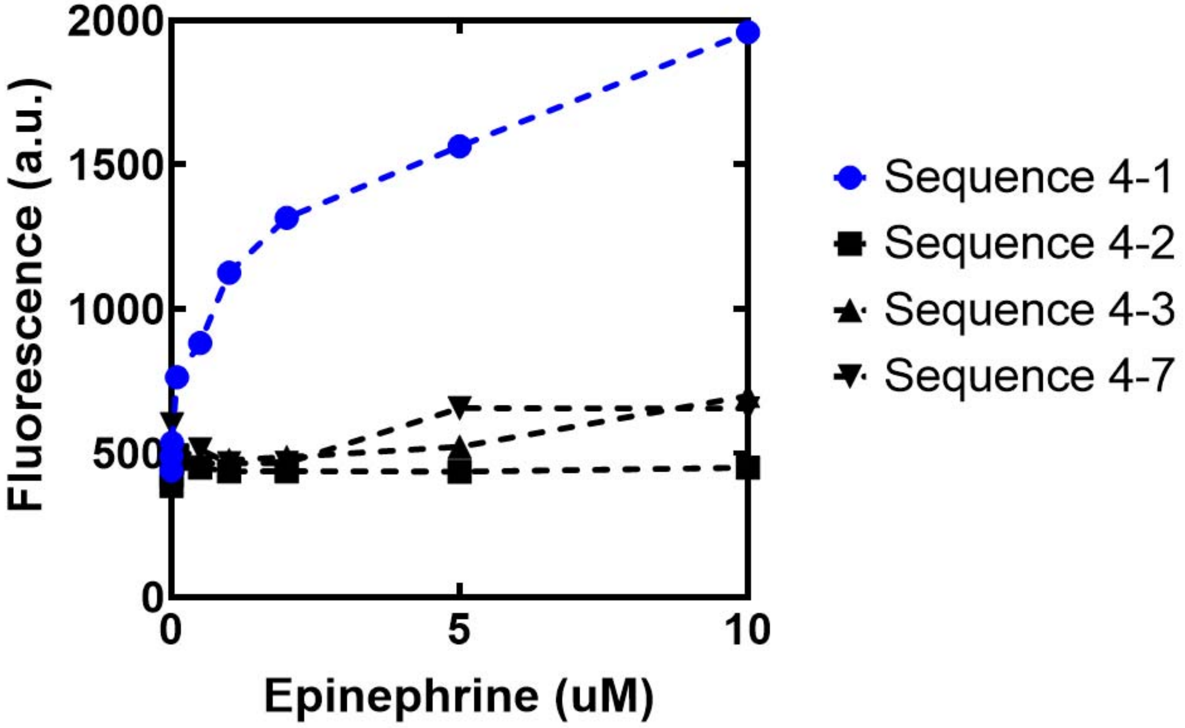

**Figure S11:** Chosen aptamer sequences for epinephrine (**Table S2**) were tested by flow cytometry assay for binding from 0 to 10 μM FITC epinephrine. $K_d$ was determined to be 0.3 μM ± 0.18 for **4-1**, and could not be determined for the other sequences.

**Table S2.**

**Table S2. Selected sequences from high-throughput DNA sequencing of epinephrine binders.**

| Sequence | Random region only |
|---|---|
| **4-1** | GTACGTGAATCCATGGGGACGGAGAGACGGCACGAACCAA |
| **4-2** | GGAGAGGGCTCAGACCAACAGAGGGGAAATGGGGCGGTGG |
| **4-3** | GGAGAGGGCTCAGACCAACAGAGGGGAAATGGGGCGGTGA |
| **4-7** | GTACGCGAATCCATGGGGACGGAGAGACGGCACGAACCAA |

**Table S3.**

**Table S3. Selected sequences from high-throughput DNA sequencing of Con A binders.**

| Sequence | Random region only |
|---|---|
| **1-1** | TATCATGGACTATACGGAGGTAGATCGGATATGCGAACCA |
| **2-1** | CTCCGCGGATCAATGCAGAGGATTGCAGATCCTCGACATG |
| **2-2** | CTTCGCGGATCAATGCAGAGGATTGCAGATCCTCAACATG |
| **3-1** | GTTGCATCTGCACGACTGGTGAGCTTGAGTGGCAGAAGAA |
| **3-2** | GTTGCATCTGCACTACTGGTGAACTTGAGTGGCAGAAGAA |
| **3-3** | GTTGCATCTGCACGACTGGTGAACTTGAGTGGCAGAAGAA |
| **4-1** | AGCGATAGGTGCACTGGGGTCCTCTAAGCGCGTTAACGAG |
| **5-1** | TAGTACGGAGGAACGTGCGAGCGGTAGCATTATAGCGAGA |
| **6-1** | CACGTACTGCTACGGGGGAGGGAGGTATCTGTCGCGGA |
| **6-2** | CACGTACTGCTACGGGAAGGGAGGTATCTGTCGCGGA |
| **7-1** | TCTGTGACGGTACGTCGCTGGAAGAAGTTGGGACGTATTA |
| **9-1** | GAAGCAAGTTGGTCTTTAACGATACAACAGCTTGCGGAAC |
| **11-1** | GGAGGTGTTACTGGCCGGGGAAGATTGAGGGTGGCGTGG |
| **17-1** | GTTGAATCTGGATACGATTTCTGAGTTCTTAATGGGAAGA |

**Figure S12.**

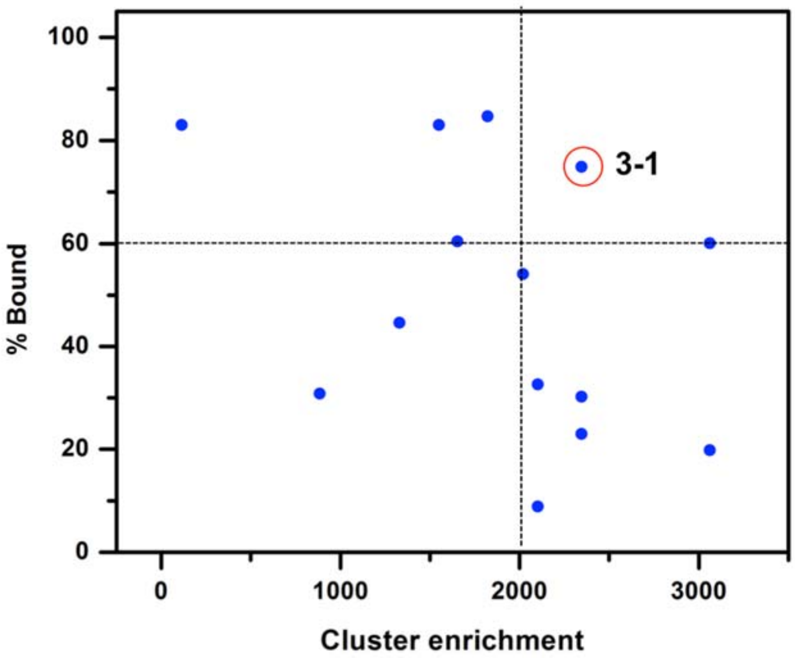

**Figure S12. Binding of selected sequences to fluorescently-labeled Con A in a particle-based assay.** Two criteria were considered to identify the top-performing base-modified aptamer. First, >60% of base-modified aptamer-displaying particles should bind Con A in a particle-based fluorescent assay. Second, the base-modified aptamer should originate from a cluster that has undergone >2,000-fold enrichment. **ConA-3-1** met both of these criteria.

**Figure S13.**

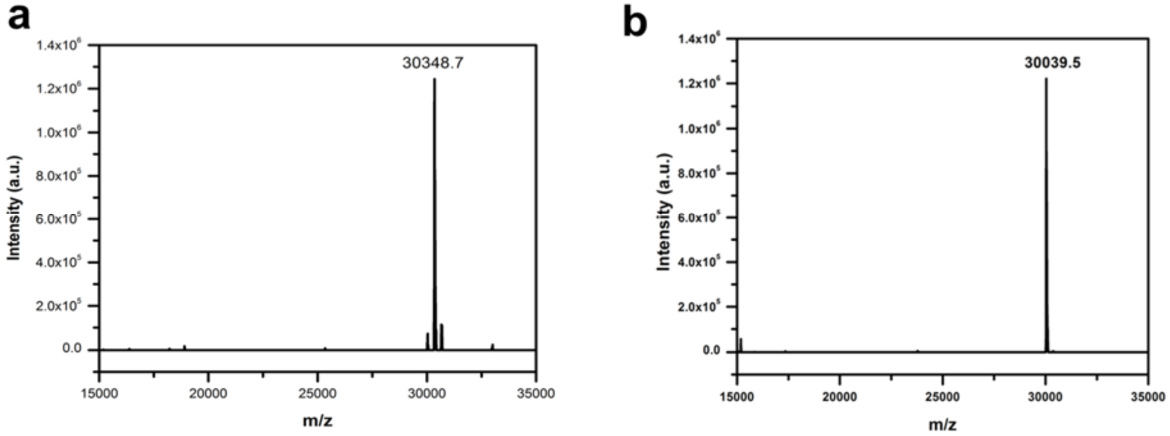

**Figure S13. ESI-MS characterization of solution-phase ConA-3-1 and ConA-3-1m with 5'-biotinylation. (a) ConA-3-1.** Expected mass: 30352.6; observed mass: 30348.7. (b) **ConA-3-1m**. Expected mass: 30040.5; observed mass: 30039.5.

**Figure S14.**

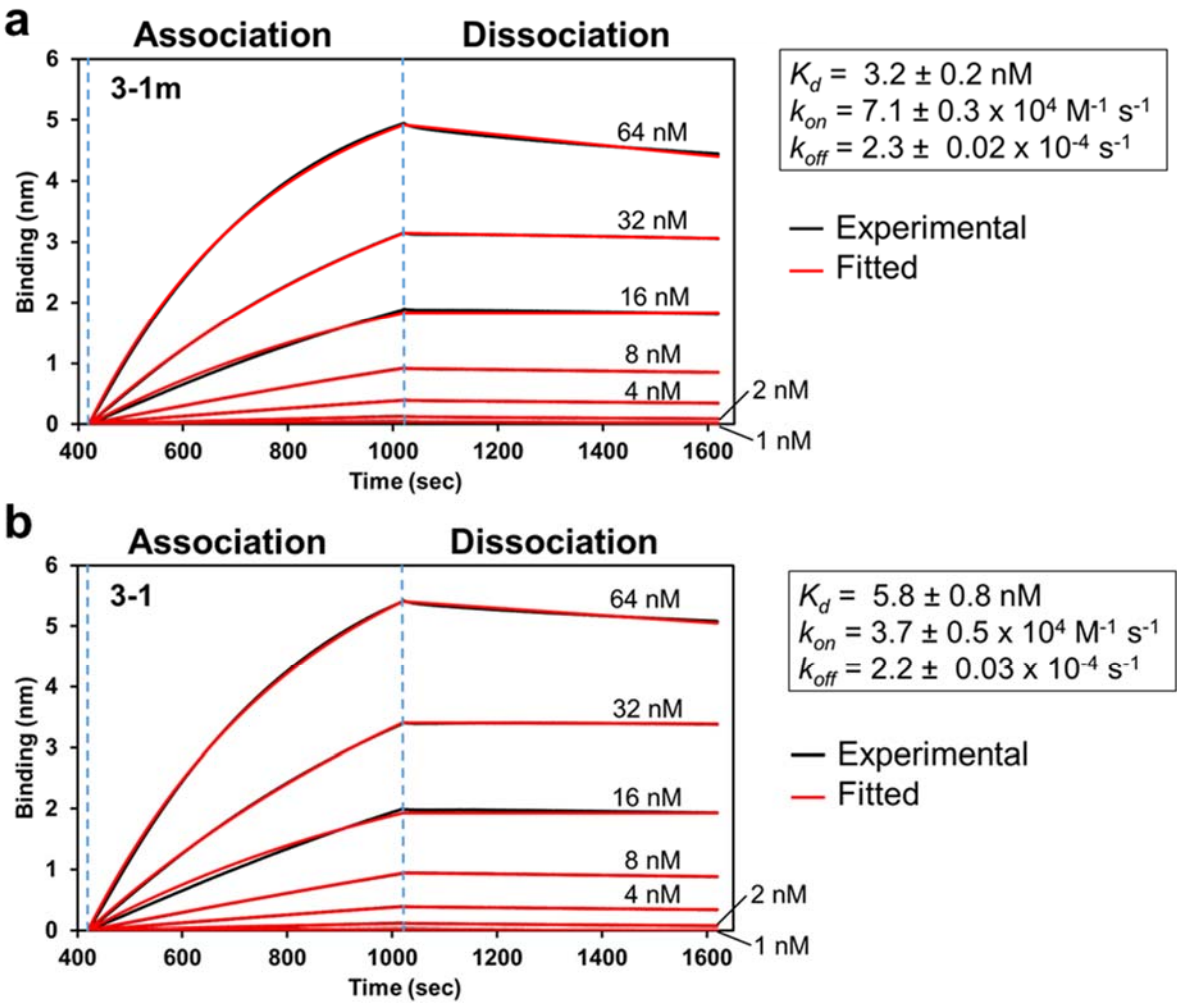

**Figure S14. BLI analysis of ConA-3-1m and ConA-3-1.** BLI measurement of Con A interacting with surface-immobilized (**a**) **ConA-3-1m** and (**b**) **ConA-3-1**. Global fitting of target association and dissociation at each concentration was performed to generate $K_d$, $k_{on}$, and $k_{off}$ values.

**Figure S15.**

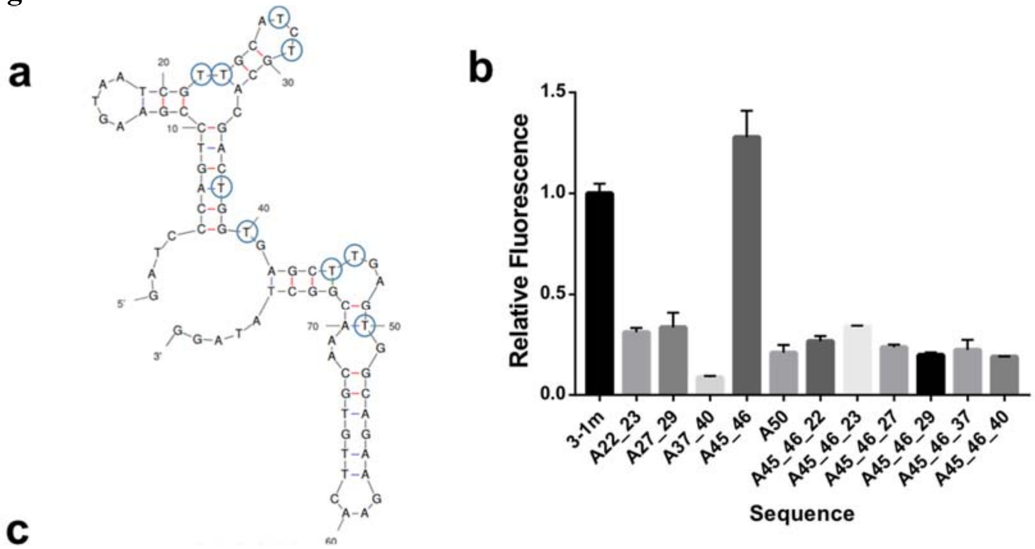

**c**

| | |
|---|---|
| **3-1m:** | GATCCCAGTCCGAAGTAATCG44GCA4C4GCACGAC4GG4GAGC44GAG4GGCAGAAGAAC44G4GCAAACGGC4A4AGG |
| **A22_23:** | GATCCCAGTCCGAAGTAATCGAAGCA4C4GCACGAC4GG4GAGC44GAG4GGCAGAAGAAC44G4GCAAACGGC4A4AGG |
| **A27_29:** | GATCCCAGTCCGAAGTAATCG44GCAACAGCACGAC4GG4GAGC44GAG4GGCAGAAGAAC44G4GCAAACGGC4A4AGG |
| **A37_40:** | GATCCCAGTCCGAAGTAATCG44GCA4C4GCACGACAGGAGAGC44GAG4GGCAGAAGAAC44G4GCAAACGGC4A4AGG |
| **A45_46:** | GATCCCAGTCCGAAGTAATCG44GCA4C4GCACGAC4GG4GAGCAAGAG4GGCAGAAGAAC44G4GCAAACGGC4A4AGG |
| **A50:** | GATCCCAGTCCGAAGTAATCG44GCA4C4GCACGAC4GG4GAGC44GAGAGGCAGAAGAAC44G4GCAAACGGC4A4AGG |
| **A45_46_22:** | GATCCCAGTCCGAAGTAATCGA4GCA4C4GCACGAC4GG4GAGCAAGAG4GGCAGAAGAAC44G4GCAAACGGC4A4AGG |
| **A45_46_23:** | GATCCCAGTCCGAAGTAATCG4AGCA4C4GCACGAC4GG4GAGCAAGAG4GGCAGAAGAAC44G4GCAAACGGC4A4AGG |
| **A45_46_27:** | GATCCCAGTCCGAAGTAATCG44GCAAC4GCACGAC4GG4GAGCAAGAG4GGCAGAAGAAC44G4GCAAACGGC4A4AGG |
| **A45_46_29:** | GATCCCAGTCCGAAGTAATCG44GCA4CAGCACGAC4GG4GAGCAAGAG4GGCAGAAGAAC44G4GCAAACGGC4A4AGG |
| **A45_46_37:** | GATCCCAGTCCGAAGTAATCG44GCA4C4GCACGACAGG4GAGCAAGAG4GGCAGAAGAAC44G4GCAAACGGC4A4AGG |
| **A45_46_40:** | GATCCCAGTCCGAAGTAATCG44GCA4C4GCACGAC4GGAGAGCAAGAG4GGCAGAAGAAC44G4GCAAACGGC4A4AGG |

**Figure S15. Structure-activity relationship of ConA-3-1m.** (a) Folding structure of **ConA-3-1m** predicted by mFold. Note that modified nucleotide **4** has been substituted with dT in the simulation. The circled nucleotide positions were mutated to dA individually or in pairs, and the binding of the mutant base-modified aptamers was characterized in a particle-based fluorescent assay. (b) The relative fluorescence signals of the mutant sequences, which are shown in (c). The error bars were derived from three experimental replicates. The fluorescence signals were first normalized to particle coating, and then to the relative signal of **ConA-3-1m**.

**Figure S16.**

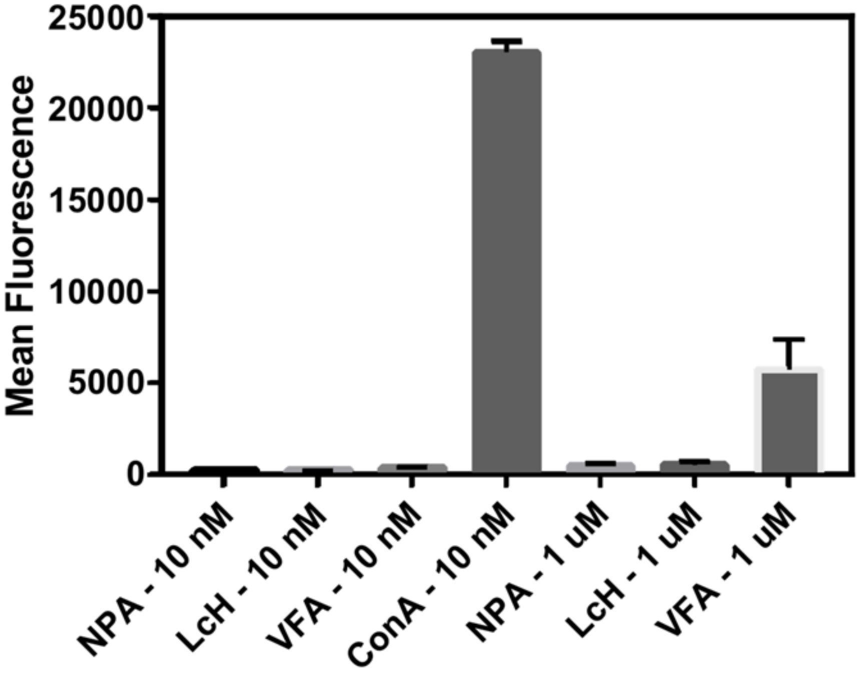

**Figure S16. ConA-3-1m exhibits little binding to NPA, LcH, and VFA lectins.** We incubated particles coated with **ConA-3-1m** with fluorescently-labeled mannose-binding lectins. These were then washed and analyzed by FACS based on mean fluorescence of the population. Error bars were derived from three experimental replicates.

Figure S17.

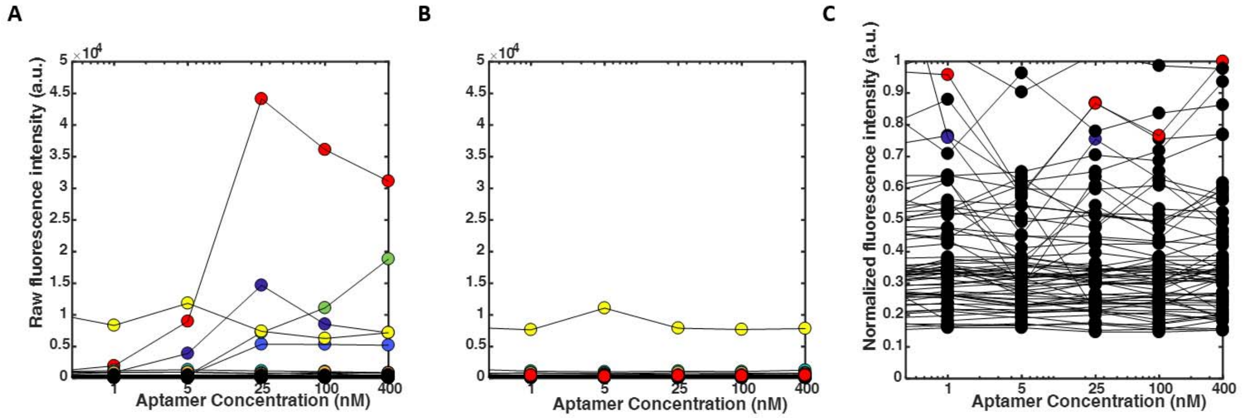

**Figure S17. Aptamer array binding for ConA-3-1m and previously described aptamer Seq 1**. (A) Raw binding signal of **ConA-3-1m** aptamer binding to Con A (red) and other lectins. We did not test concentrations high enough to determine a $K_d$ value for off-target binding. The normalized plot of this data is shown as **Fig 6B**. (B) Raw binding signal of the previously published **Seq 1** aptamer (red) to Con A and other lectins on an array, scaled to match the axes for the raw binding signal of **ConA-3-1m**. The signal produced by **Seq 1** is negligible compared to our aptamer, and only VVA (yellow) continues to show non-specific binding. (C) Normalized signal of **Seq 1** binding to Con A (red) and other lectins. The signal from each lectin was normalized to the maximum Con A signal measured in each assay.

**Table S4.**

**Table S4. Additional information on lectins spotted on the lectin array.**

The following information is replicated from Lectin Array 70 product manual.

#### VI. Lectin Array 70 Key

| | Lectins | Abbreviation | Source | Carbonhydrate specificity |
|---|---|---|---|---|
| 1 | *Anguilla anguilla* | AAA | *Anguilla anguilla* (Fresh Water Eel) | αFuc |
| 2 | *Aleuria aurantia* | AAL | *Aleuria aurantia* mushrooms | Fucα6GlcNAc |
| 3 | *Agrocybe cylindracea lectin* | ACG | E. coli expressed *Agrocybe cylindracea* galectin | α2-3 Sialic Acid |
| 4 | *Amaranthus caudatus* | ACL, ACA | *Amaranthus caudatus* seeds | Galβ3GalNAc |
| 5 | *Allium sativum* | ASA | *Allium sativum* agglutinin (Garlic) | αMan |
| 6 | *Musa acuminata lectin* | BanLec | E. coli expressed *Musa acuminata* | Mannose, Glucose, branched high-mannose containing α1,3-glycoside bond |
| 7 | *Burkholderia cenocepacia lectin* | BC2L-A | E. coli expressed *Burkholderia cenocepacia* | High-mannose |
| 8 | *Burkholderia cenocepacia lectin* | BC2LCN (AiLecS1) | E. coli expressed *Burkholderia cenocepacia* | Fucα1-2Galβ1-3GalNAc (H type 3), Fucα1-2Galβ1-3GlcNAc (H type 1) |
| 9 | *Bauhinia purpurea* | BPA, BLP | *Bauhinia purpurea alba* (Camel's Foot Tree) | Galβ3GalNAc |
| 10 | *Calystegia sepium lectin* | Calsepa | E. coli expressed Calystegia sepium | High-mannose |
| 11 | *Coprinopsis cinerea lectin* | CGL2 | E. coli expressed *Coprinopsis cinerea* | βGal, GalNAcα1-3Gal (Blood Group A), Galα1-3Gal (Blood Group B) |
| 12 | *Clitocybe nebularis lectin* | CNL | E. coli expressed *Clitocybe nebularis* | α/βGalNAc, GalNAcβ1-4GlcNAc, GalNAcα1-3[Fucα1-2]Galβ1-4GlcNAc (Blood Group A) |
| 13 | Coanavalin A | Con A | *Coanavalia ensformis* (Jack Beans) seeds | αMan, αGlc |
| 14 | *Dolichos biflorus* | DBA | *Dolichos biflorus* (Horse Gram) seeds | αGalNAc |
| 15 | *Dictyostelium discoideum lectin* | Discoidin I | E. coli expressed *Dictyostelium discoideum* | αGalNAc (Tn antigen), LacNAc |
| 16 | *Dictyostelium discoideum lectin* | Discoidin II | E. coli expressed *Dictyostelium discoideum* | Gal, LacNAc, Asialoglycans, Gal/GalNAcβ1-4GlcNAcβ1-6Gal/GalNAc |
| 17 | *Datura stramonium* | DSA, DSL | seeds | $(GlcNAc)_{2-4}$ |
| 18 | *Erythrina cristagalli* | ECA, ECL | *Erythrina cristagalli* (Coral Tree) seeds | Galβ4GlcNAc |
| 19 | *Eunonymus europaeus* | EEL | *Eunonymus europaeus* (Spindle Tree) seeds | Galα3Gal |
| 20 | *E. coli lectin* | F17AG | E. coli expressed *E. coli* | GlcNAc |
| 21 | Human galectin1 lectin (stable form) | Gal1 | E. coli expressed human galectin1 (stable form) | branched LacNAc, Gal |

14

| | Lectins | Abbreviation | Source | Carbonhydrate specificity |
|---|---|---|---|---|
| 22 | Human galectin1-S lectin | Gal1-S | E. coli expressed human galectin1-S | branched LacNAc |
| 23 | Human galectin2 lectin | Gal2 | E. coli expressed human galectin2 | GalNAcα1-3Gal (Blood Group A), branched LacNAc |
| 24 | Human galectin3 lectin (full-length) | Gal3 | E. coli expressed Human galectin3(full-length) | poly LacNAc |
| 25 | Human galectin 3C-S lectin | Gal3C-S | E. coli expressed Human galectin 3C-S | poly LacNAc |
| 26 | Human galectin7-S lectin | Gal7-S | E. coli expressed Human galectin7-S | Galβ1-3GlcNAc |
| 27 | Human galectin9 lectin (Stable Form) | Gal9 | E. coli expressed human galectin9 | poly LacNAc, GalNAcα1-3Gal (Blood Group A) |
| 28 | *Galanthus nivalis* | GNA, GNL | *Galanthus nivalis* (Snowdrop) bulbs | αMan |
| 29 | *Griffithia sp. Lectin* | GRFT | E. coli expressed *Griffithia sp.* | High-mannose |
| 30 | *Griffonia (Banderaea) simplicifolia I* | GS-I, GSL-II, BSL-I | *Griffonia (Banderaea) simplicifolia* seeds | αGal, α3GalNAc |
| 31 | *Griffonia (Brandeiraea)* | GS-II, GSL-II, BSL-II | *Griffonia (Banderaea) simplicifolia* seeds | α or βGlcNAc |
| 32 | *Hippeastrum hybrid* | HHA, HHL, AL | *Hippeastrum* hybrid (Amaryllis) bulbs | αMan |
| 33 | Jacalin | Jacalin, AIL | *Artocarpus integrifolia* (Jackfruit) seeds | Galβ3GalNAc |
| 34 | *Phaseolus lunatus* | LBA | *Phaseolus lunatus* (Lima Bean) seeds | GalNAcα(1,3)[αFuc(1,2]Gal |
| 35 | *Lens Culinaris* | LcH, LCA | *Lens culinaris* (lenti l) seeds | αMan, αGlc |
| 36 | *Lycopersicon esculentum* | LEA, LEL, TL | *Lycopersicon esculentum* (tomato) fruit | $(GlcNAc)_{2-4}$ |
| 37 | Lentil lectin | Lentil | *Lens culinaris* seeds | D-Mannose, D-glucose |
| 38 | *Lotus tetragonolobus* | Lotus, LTL | *Lotus tetragonolobus, Tetragonolobus purprea(* Winged Pea, Asparagus Pea) seeds | αFuc |
| 39 | *Laetiporus sulphureus lectin* | LSL-N | E. coli expressed *Laetiporus sulphureus* | LacNAc, poly LacNAc |
| 40 | *Maackia amurensis I* | MAA, MAL, MAL-I | *Maackia amurensis* seeds | Galβ4GlcNAc |
| 41 | Human malectin lectin | Malectin | E. coli expressed human malectin | $Glc_2$-*N*-biose |
| 42 | *Marasmius oreades lectin* | MOA | E. coli expressed *Marasmius oreades* | Galα1-3[Fucα1-2]Galβ1-4GlcNAc (Blood Group B) Galα1-3Galβ1-4GlcNAc, Galα1-3Gal |
| 43 | *Maclura pomifera* | MPL, MPA | *Maclura pomifera* (Osage Orange) seeds | Galβ3GalNAc |
| 44 | *Narcissus pseudonarcissus* | NPA, NPL, DL | *Narcissus pseudonarcissus* (Daffodil) bulbs | αMan |
| 45 | *Oryza sative lectin* | Orysata | E. coli expressed *Oryza sative* | High-mannose |
| 46 | *Pseudomonas aeruginosa lectin* | PA-IIL | E. coli expressed *Pseudomonas aeruginosa* | Fucose, Fucose containing oligosaccharides, Mannose |
| 47 | *Pseudomonas aeruginosa lectin* | PA-IL | E. coli expressed *Pseudomonas aeruginosa* | Galα1-3(4)Gal |
| 48 | *Phlebodium aureum lectin* | PALa | E. coli expressed *Phlebodium aureum* | High-mannose |
| 49 | *Phaseolus vulgaris Erythroagglutinin* | PHA-E | *Phaseolus vulgaris Erythroagglutinin* (Red Kidney Bean) seeds) | Galβ4GlcNAcβ2Manα6(GlcNAcb4)(GlcNAcβ4Manα3)Manβ4 |
| 50 | *Leucoagglutinin* | PHA-L | *Phaseolus vulgaris Erythroagglutinin* (Red Kidney Bean) seeds) | Galβ4GlcNAcβ6(Gl cNAcβ2Manα3)Manα3 |
| 51 | Phaseolus vulgaris agglutinin | PHA-P | *Phaseolus vulgaris Erythroagglutinin* (Red Kidney Bean) seeds) | Galβ4GlcNAcβ2Manα6(GlcNAcb4)(GlcNAcβ4Manα3)Manβ4, Galβ4GlcNAcβ6(Gl cNAcβ2Manα3)Manα3 |
| 52 | Peanut | PNA | Arachis hypogaea *Peanut* | Galβ3GalNAc |
| 53 | *Pleurocybella porrigens lectin* | PPL | E. coli expressed Pleurocybella porrigens | α/βGalNAc |
| 54 | *Pisum sativum* | PSA, PEA | *Pisum sativum* (Pea) seeds | αMan, αGlc |
| 55 | *Polyporus squamosus lectin* | PSL1a | E. coli expressed *Polyporus squamosus* | α2-6 Sialic Acid |
| 56 | *Psophocarpus* | PTL, PTL-I, WBA-I | *Psophocarpus tretragonolobus* (Winged Bean) | GalNAc, Gal |
| 57 | *Ralstonia solanacearum lectin* | RS-Fuc | E. coli expressed *Ralstonia solanacearum* | Fucose |
| 58 | Sambucus Sieboldiana Lectin | SAMB | Japanese elderberry | NeuAcα2-6Gal/GalNAc |
| 59 | Soybean | SBA | *Glycine max* (Soybean) seeds | α > βGalNAc |
| 60 | *Sophora japonica* | SJA | *Sophora japonica* (Japanese Pagoda Tree) seeds | βGalNAc |
| 61 | *Sambucus nigra I* | SNA-I | *Sambucus nigra* (Elderberry) bark | NANAα(2,6)GalNAc > GalNAc = Lac > GalNANAα(2,6)Gal |
| 62 | *Sambucus nigra II* | SNA-II | *Sambucus nigra* (Elderberry) bark | GalNAc > Gal |
| 63 | *Solanum tuberosum* | STL, PL | *Solanum tuberosum,* (potato) tubers | $(GlcNAc)_{2-4}$ |
| 64 | *Urtica dioica* | UDA | *Urtica dioica* (Stinging Nettle) seeds | GlcNAc |
| 65 | *Ulex europaeus I* | UEA-I | *Ulex europaeus* (Furze Gorse) seeds | αFuc |
| 66 | *Ulex europaeus II* | UEA-II | *Ulex europaeus* (Furze Gorse) seeds | Poly β(1,4)Gl cNAc |
| 67 | *Vicia faba* | VFA | *Vicia faba* (Fava Bean) seeds | αMan |
| 68 | *Vicia villosa* | VVA, VVL | *Vicia villosa* (Hairy Vetch) seeds | GalNAc |
| 69 | *Wisteria floribunda* | WFA | *Wisteria floribunda* (Japanese Wisteria) seeds | GalNAc |
| 70 | Wheat Germ | WGA | *Triticum volganis* (Wheat Germ) | GlCNAc |

| Sugar Abbreviations | | | |
|---|---|---|---|
| Fuc: L-Fucose | Gal: D-Galactose | GalNAc: *N*-Acetylglactosamine | Glc: D-Glucose |
| GlcNAc: *N*-Acetylglucosamine | Lac: Lactose | Man: Mannose | |
